# Revealing Free Energy Landscape from MD Data via Conditional Angle Partition Tree

**DOI:** 10.1101/2021.09.27.461919

**Authors:** Hangjin Jiang, Han Li, Wing Hung Wong, Xiaodan Fan

## Abstract

Deciphering the free energy landscape of biomolecular structure space is crucial for understanding many complex molecular processes, such as protein-protein interaction, RNA folding, and protein folding. A major source of current dynamic structure data is Molecular Dynamics (MD) simulations. Several methods have been proposed to investigate the free energy landscape from MD data, but all of them rely on the assumption that kinetic similarity is associated with global geometric similarity, which may lead to unsatisfactory results. In this paper, we proposed a new method called Conditional Angle Partition Tree to reveal the hierarchical free energy landscape by correlating local geometric similarity with kinetic similarity. Its application on the benchmark alanine dipeptide MD data showed a much better performance than existing methods in exploring and understanding the free energy landscape. We also applied it to the MD data of Villin HP35. Our results are more reasonable on various aspects than those from other methods and very informative on the hierarchical structure of its energy landscape.

## 1 Introduction

Proteins and nucleic acids depend on their biomolecular structures to perform their biological functions properly. These biomolecules may also undergo reversible transitions between alternative structures called different conformations. Conformations have different amounts of Gibbs free energy, which characterize their different thermodynamic stability, with lower free energy corresponding to more stable status. The free energy landscape of a conformational space is usually rugged [1], [2], with a number of high-energy barriers partitioning the space into a set of metastable low-energy wells, which are also called macrostates [3]. It is crucial to understand the free energy landscape and the conformational dynamics within and between energy wells in order to study many biological or chemical processes, such as protein-protein interaction, RNA fold, and protein folding. Improper conformational changes, such as protein misfolding, are linked to many serious diseases, such as Alzheimer’s disease, Bovine spongiform encephalopathy, Huntington’s disease, Parkinson’s disease, cancers and cancer-related syndromes [4].

Several existing experimental techniques can be used to study conformational changes at the atomic level to certain extent. For example, X-ray imaging can provide static snapshots of biomolecules; nuclear magnetic resonance can provide dynamic information for small biomolecular systems; single molecule fluorescence resonance energy transfer can provide dynamic information on conformational changes along certain order parameter such as the distance between a probe pair; Cryo-electron microscopy can determine biomolecular structures at near-atomic resolution. However, it is either impossible or too expensive for these experimental techniques to provide global dynamic information on conformational changes at atomic resolution. To complement this shortage, Molecular Dynamics (MD) and Monte Carlo computer algorithms are used to simulate conformational trajectories [5], [6], [7].

Molecular Dynamics (MD) simulations sample from a conformational space by producing a number of trajectories. Each trajectory records a sequence of conformations at times *t* = 0, *τ*, 2*τ*, …, *nτ*, where *τ* denotes the observation interval. Due to the complexity of the rugged free energy landscape, generalized ensemble algorithms, such as multicanonical algorithm [8], Replica Exchange [9] and long time dynamical simulations [10], [11], are used in MD simulations to generate a wider sampling by helping the simulation trajectories pass through energy barriers with a higher probability and avoid trapping in local modes [12].

However, there is a gap between the timescale of computer simulations and the timescale of typical real biological conformational changes. Since the conformational space is a high-dimensional continuous coordinate space, the raw simulation trajectories may contain thousands of conformations, but very few transitions between any specific pair of these conformations can be observed from the simulation trajectories, i.e., the entries of transition matrix between conformations are almost zero. The aim of analyzing MD data is to find the metastable structures of the biomolecular and study the dynamics between them. To elucidate the free energy landscape of the conformational space from these simulated trajectories, a two-step procedure [3], [13], [14], [15] is commonly used, such as Perron Cluster Cluster Analysis (PCCA) and its variants [3], [16], [17], Most Probable Path (MPP) [13], Gibbs sampling [14], Automatic Partitioning for Multibody systems (APM) [18], transition disconnectivity graph (TRDG) [19], [20] and Minimum Variance Clustering Approach (MVCA) [21].

Typically, a two-step procedure is used to learn metastable states from MD data. In the first step, which is called the splitting step, conformations are grouped into a number of small tight sets called microstates according to their geometric similarity. This is the so called geometric clustering step, which is a kind of clustering for vectorial data [22]. With a high similarity threshold, the conformations within a microstate can have both geometric and kinetic similarity, which ensures fast converting among them. We say two conformations are geometrically similar, if they are close in geometric distance. Kinetic (dynamic) similarity between two conformations (microstates) is measured by the transition probability between them. Higher transition probability implies more similar in dynamics. The transition probability matrix between two conformations (microstates) is obtained by normalizing the transition matrix such that the sum of each row is 1, see Supplementary Figure 1 for an illustration. After mapping simulation trajectories from a sequence of conformations to a sequence of microstates and making use of the time-reversible/detailed-balance nature of MD simulations [23], the connections among all microstates are used to form a transition matrix, where each cell denotes the counts of transitioning from its column microstate to row microstate [24].

The second step, which is called the lumping step, is to cluster microstates into macrostates based on the transition matrix beween microstates at the microstate level. This is the so-called kinetic (dynamic) clustering step. It is a special type of clustering on pairwise similarity data or weighted network data [25]. Several classes of methods have been proposed for this problem. One class of methods is based on main eigenvectors of the transition matrix, such as PCCA [26] and its modified versions [3], [17], [24]. Another class of methods formulate the problem as a steepest-descent-search or minimum-cut problem based on the idea of maximizing the sum of intra-macrostate transition probabilities, such as the original steepest descent method by [27], the iterative K-medoid partitioning and lumping algorithm [16], the cut-based free-energy profile [28] and MPP [13]. Except for these specialized methods, some general clustering methods for pairwise similarity data may also, in principle, be adapted to this kinetic clustering problem, such as k-means clustering [29], self-organizing maps [30], maximum entropy [31] and deterministic annealing [32]. But all these methods can neither efficiently handle the sparseness of the microstate transition matrix nor provide a comprehensive hierarchical view of the number of clusters and the cluster structure. For example, the methods based on eigenvector calculation tend to be numerically unstable due to the sparse counts resulting from insufficient sampling [24]. Also, the number of clusters is assumed known for most of existing clustering methods, with few exceptions such as the Chinese restaurant process approach [33], [34], which is only applicable to vector data. Therefore, they cannot be applied to the energy landscape case due to its multi-resolution nature, i.e., the number of clusters essentially depends on which resolution the algorithm is sampling in the energy hierarchy.

In the above two-step procedure, the first step only uses the geometric similarity information while the second only uses the kinetic similarity. See [35] for more discussion on this procedure. In this paper, we proposed a new method, CAPT, which makes use of both geometric and kinetic similarity information throughout the whole process. Our method is built on the following facts: 1) local structures, more specifically the dihedral angles of amino acid residues in the backbone is the main source of the energy wells [36], [37]; 2) the distribution of dihedral angles contributing to the free energy landscape is not uniform. These facts make the joint space of dihedral angles highly multi-modal, thus forming different energy wells. Our new method selects important backbone dihedral angles to partition the joint space of all dihedral angles in a tree structure. We apply our method to two molecular systems to show that our method can elucidate the conformational space efficiently.

## 2 Methods

Our method is motivated by two key observations. The first one is from the cause of energy barriers. The second one comes from the separated use of geometrical similarity and dynamical similarity in the two-step procedure taken by most existing methods, as described previously. We shall elaborate more on these two observations.

### 2.1 Energy Barrier

The first key observation is about the basic knowledge of biomolecular energy wells. The existence of energy barriers is due to the minimum distance between two atoms. For example, the minimum distance between two *H* atoms is 2.0 Å, and that between an *H* and a *C* atom is 2.4 Å [38]. The distance between two atoms changes when related residues rotate. If the distance between two atoms becomes closer to their minimum distance, it becomes more difficult for this residue to rotate further in the same direction. This explains why the range of rotation of this residue is restricted to a subregion/interval of [−*π, π*] in the Ramachandran plot (also called Ramachandran diagram or [*ϕ, ψ*] plot) of biomolecules. Importantly, the evolution between different conformations is mainly caused by such kind of rotations [38], [39], [40], and the backbone torsion angles are used to build native protein conformations [41], which implies the torsion angles are informative to protein structures. Furthermore, it was shown that normal modes in torsion angle space better correlate with conformation changes in proteins [42]. Thus, we use backbone torsion angles to describe biomolecules. This strategy is also used in, for example, [36], [37], [43], [44].

### 2.2 Analysis of the Two-step Procedure

We secondly observe that the two-step procedure with splitting step and lumping step is taken by most popular methods, such as PCCA [16], [45], PCCA+ [17], [46], MPP [13], and Gibbs sampling method [14]. The splitting step clusters conformations into microstates using K-means/K-medoids based on principal components of 3-D coordinate data or backbone torsion angle data. The lumping step clusters microstates into macrostates. Essentially, the splitting step clusters conformations into microstates based on geometrical similarity, while the lumping step lumps microstates into macrostates based on dynamical similarity.

The key assumption for the splitting step is that two conformations are in the same microstate if they are geometrically similar. PCCA, PCCA+ and MPP use K-means/K-medoids in this step to cluster conformations into microstates. However, one can not make sure results obtained by K-means/K-medoids have the following clustering property: conformations belonging to the same microstate are around a small neighborhood of the center of this microstate. More specifically, we shall require an *ε*-cover of the whole space with centers {ℙ_1_, ℙ_2_, …. ℙ_*n*_} such that for each conformation ℙ, there is an ℙ_*i*_, such that *d*(ℙ, ℙ_*i*_) ≤ *ε*, where *d*(·, ·) is a distance function, and it can be defined on principal components of torsion angles, see for example, [13], [43], [44], [47]. There are two difficulties for constructing such an *ε*-cover. Firstly, it is hard, especially in high dimensional space, to make sure balls with center *P*_*i*_ and radius *ε* are disjoint. Secondly, it is difficult to determine *ε*. If *ε* is too small, there are too many microstates to handle with, and the transition frequency between two microstates will be very low. If *ε* is too large, the cluster structure will be covered, which induces additional difficulty in exploring the landscape. Furthermore, due to the huge number of conformations to be clustered, the K-means/K-medoids algorithm usually converges to a local mode instead of a global mode.

In the lumping step, based on microstates found in the splitting step, both PCCA and PCCA+ try to group the microstates into macrostates by maximizing the sum of diagonals of the transition matrix among macrostates [16], [17], [45]. For MPP, this dynamical clustering is based on the so called most probable pathway. More specifically, MPP merges a microstate with its neighboring microstate on its most probable pathway which has the lowest free energy. Basically, PCCA, PCCA+ and MPP do not consider geometrical similarity between conformations in this step, and fully depends on the dynamical information. Different from these methods, Gibbs sampling algorithm tries to cluster microstates into macrostates based on a Bayesian Poisson model, which also depends only on the dynamical information.

Methods mentioned above consider separately the geometrical similarity and dynamical similarity, which is not reasonable. Undoubtedly, conformations in the same microstate are of both high geometrical similarity and high dynamical similarity. For macrostates, the dynamical similarity may come from their local geometrical similarity. However, this kind of relationship between geometrical similarity and dynamical similarity is ignored by other methods such as MPP, APM and PCCA/PCCA+. This motivated us to always consider both types of similarity simultaneously.

### 2.3 Definitions

In the following, we give definitions of key elements for our algorithm: partition score, mean absolute distance and one-step distance

#### Definition 1

(Partition score of angle *θ*). *Assume that we have observations* 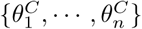 *for a given dihedral angle θ in parent cluster C, where n is the number of frames/conformations in the cluster C. If there are k local modes in the density of θ, we partition frames in this cluster into k sub-clusters (states) by naive Bayesian classifier [48]. Then the original MD trajectories of conformations are transformed into trajectories of these k states, and we obtain a k* × *k transition probability matrix A between these states. We score this angle θ by* min_*i*_ *A*(*i, i*), *i.e*., *the minimum self-transition probability*.

#### Algorithm 1

Conditional Angle Partition Tree (CAPT)

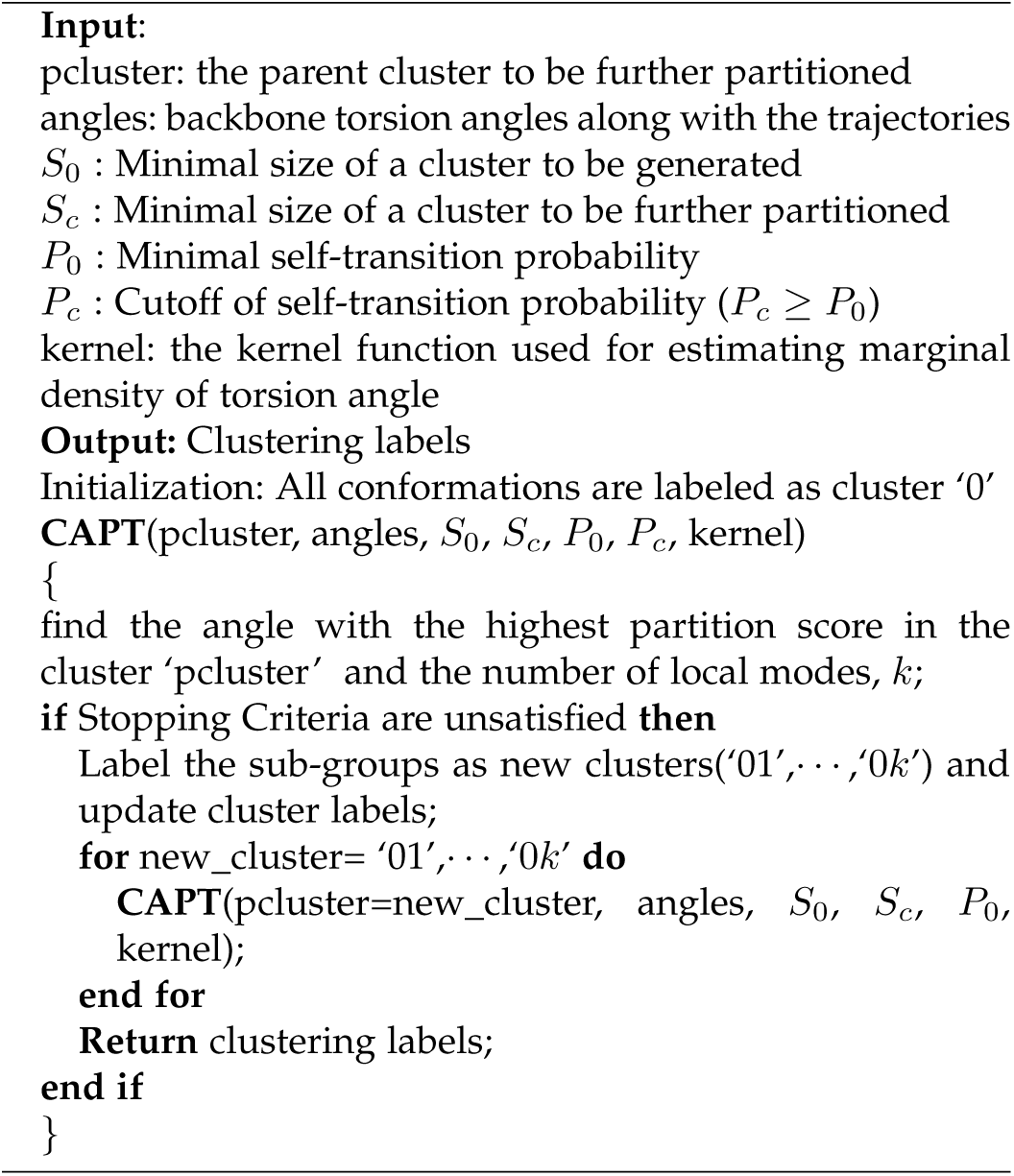

#### Definition 2

(Mean Absolute Distance). *The Mean Absolute Distance (MAD) between two conformations, P*_*i*_ = (*θ*_*i*1_, *θ*_*i*2_, · · ·, *θ*_*ik*_) *for i* = 1, 2, *is defined by*

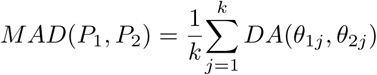

*where θ*_*ij*_ *is a torsion angle, DA*(*θ*_1*j*_, *θ*_2*j*_) *is the distance between two torsion angles. Since the torsion angles are periodic with period* 2*π, the distance between two torsion angles (DA) is defined by DA*(*θ*_1_, *θ*_2_) = min{|*θ*_1_ − *θ*_2_|, |*θ*_1_ − *θ*_2_ + 2*π*|, |*θ*_2_ − *θ*_1_ + 2*π*|}.

#### Definition 3

(One-step Distance). *The one-step distance is the distance between the conformations in adjacent frames along the MD trajectory*.

Typically, the one-step distance can be one-step MAD, one-step RMSD, or one-step DA.

### 2.4 The CAPT algorithm

Finding the stable states/structures of a biomolecule based on the MD data, statistically, is to estimate its landscape density function 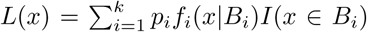, where *k* is the number of stable states, {*B*_1_, *B*_2_, *⋯, B*_*k*_} are disjoint partitions/basins of the whole conformation space Ω (i.e., 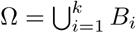), *p*_*i*_ is the probability of a conformation belonging to basin *B*_*i*_ such that 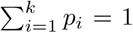, and *f*_*i*_(*x B*_*i*_) is the density function of conformations belonging to basin *B*_*i*_ such that 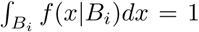. Each *B*_*i*_ corresponds to a stable state of the biomolecule, and the stable structure in each basin is the one with the highest density in it.

The task of joint partitioning and density estimation is related to the work of [49], which extended the classification tree for regression to handle density estimation. However, estimating *L*(*x*) is totally different from the traditional density estimation problem due to the special underlying property of basins that conformations within each basin are of high dynamical similarity due to local geometrical similarity, and the global similarity may not necessarily lead to dynamical similarity. This special property makes the partitions be the most important part of the landscape density function.

Based on the previous two observations, we design an algorithm called Conditional Angle Partition Tree (CAPT) to estimate the partitions, which is summarized in **Algorithm 1** and illustrated in Figure 1. Essentially, CAPT is similar to the classification tree method designed originally for regression. It goes recursively. We illustrate the basic idea of CAPT through the example of Alanine dipeptide step by step.

**Fig. 1.**
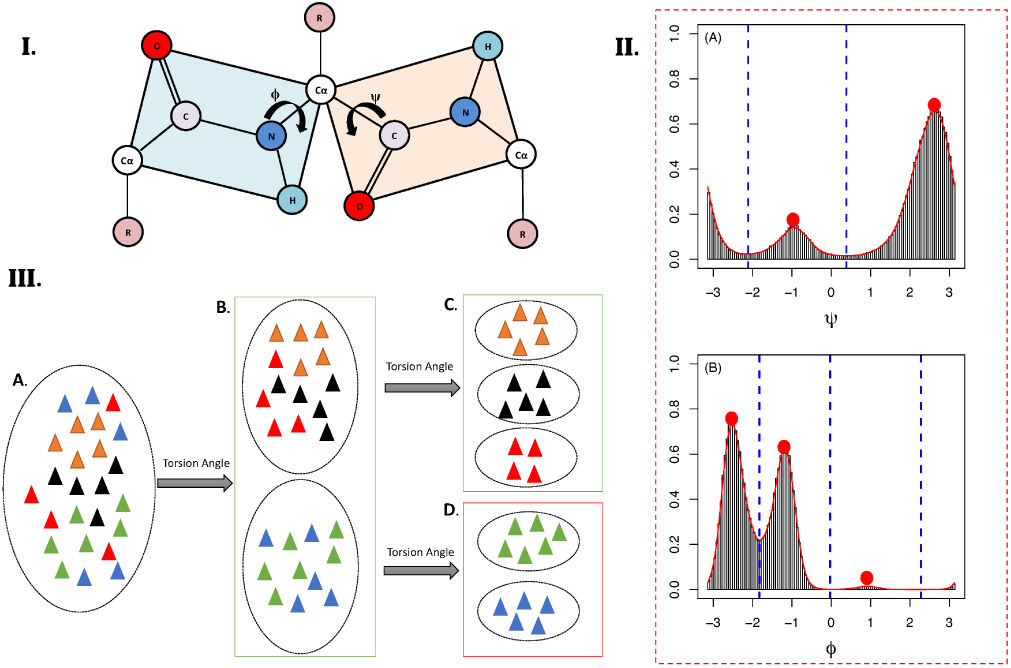
Key ideas of Conditional Angle Partition Tree (CAPT). (I) is an illustration of torsion angles (*ϕ, ψ*); (II) shows the marginal distributions of torsion angles (*ϕ, ψ*) of Alanine dipeptide summarized from its MD data. As shown in the figure, *ψ* has two local modes (denoted by red circles), and *ϕ* has three local modes. (III) shows an illustration of CAPT algorithm. The torsion angle used in each iteration of the algorithm is the one of the highest partition score within the remaining angles.

Alanine dipeptide is a small biomolecule with only two backbone torsion angles, say *ϕ* and *ψ*. The histograms with corresponding density functions of *ϕ* and *ψ* are shown in Figure 1(II), where the density functions are estimated by the kernel method [50] based on the von Mises kernel. As shown in the figure, *ψ* has two local modes (denoted by red circles), and *ϕ* has three local modes. Thus, if we use *ψ* to cluster conformations by cutting the marginal distribution of *ψ* at the point with lowest density between two local modes as shown by the blue dashed line in Figure 1(II), we get two clusters. Now, by mapping the cluster labels of conformations onto the trajectories of conformations, we obtain the transition matrix (see Supplementary Table 7) between these two clusters. The partition score of *ψ*, defined as the minimal self-transition probability, i.e., the minimal of diagonals of transition matrix, is 0.9728. Similarly, the partition score of *ϕ* is 0.9395. Thus, in this iteration, we take *ψ*, the angle with highest partition score, to partition the parent cluster (all conformations in the current iteration) into two child clusters. This is the first iteration of the CAPT algorithm for Alanine dipeptide, i.e., going from Figure 1(III.A) to Figure 1(III.B).

Next, CAPT takes each child cluster as a parent cluster and applies the clustering procedure in the above until one of the stopping criteria (see below) is meet. That is, we goes from Figure 1(III.B) to Figure 1(III.C) and (III.D). Finally, CAPT gives us a partition tree as shown in Figure 3. In summary, CAPT uses *ψ* firstly to partition all conformations into two child clusters, S01 and S02. Next, taking the new clusters as parent clusters, CAPT uses *ϕ* to partition conformations in each new parent cluster into three child clusters. We can see that conformations in different levels of nodes (clusters) are of different level of local similarity. According to our partition rule, they are also of high dynamical similarity.

### 2.5 Stopping Criteria of CAPT

CAPT has three stopping criteria. (1) All torsion angles are non-informative for clustering, that is, all torsion angles has a unimodal distribution on each terminal node. (2) It generates clusters that are not metastable, that is, the minimal self-transition probability of these clusters, min_*i*_ *p*_*ii*_, is smaller than some threshold *P*_*c*_ *> P*_0_ ≥ 0.5. Since for each *B*_*i*_, the self-transition probability *p*_*ii*_ is larger than the probability of its transition to other basins. The threshold *P*_0_ can be understood as the minimal energy barrier leading to energy wells, and *P*_*c*_ is a cutoff of the energy landscape. The suggested value for *P*_*c*_ ranges from 0.6 to 0.7, and *P*_0_ is usually set as 0.5 or just taken as *P*_*c*_. (3) It generates small meaningless clusters. To avoid meaningless small clusters, one may focus only on the basins with sample size larger than *s*_0_, equally *p*_*i*_ *> p*_0_ (see the definition of landscape density *L*(*x*)). To this aim, one may stop CAPT algorithm on a cluster with sample size smaller than a given threshold *S*_*c*_. Also, one may stop CAPT algorithm if partitioning a cluster gives at least one child cluster with sample size smaller than another threshold *S*_0_. Note that *S*_*c*_ and *S*_0_ are different, although one may set *S*_*c*_ = *S*_0_ for convenience.

### 2.6 Identifying Meaningful Metastable States and Structures

For a complex biomolecule, CAPT may produce many clusters, some of small size, some others not stable. Thus we need a method to recognize meaningful stable clusters within the partition tree. Recall that the number of conformations in each cluster reversely relates to the free energy of the cluster. That is, if a conformation is stable (with a low free energy), there should be many other geometrically similar conformations within its small neighbourhood. We thus use the local density [51], 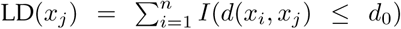, to find meaningful clusters, and consequently the stable structure and number of stable states, where *x*_*i*_ denotes the *i*-th conformation, *n* is the number of conformations, *d*_0_ is the threshold for identifying neighborhoods, and *d*(·,·) is the Mean Absolute Distance between two structures. In fact, Each frame has two kinds of local density: the local density within the cluster it belongs to, denoted by LDc, and the local density within all frames of the dataset, denoted by LDa. If the clusters are well separated, LDc equals to LDa. Thus, the difference between LDc and LDa can be used as an index for the quality of clusters.

## 3 Results

### 3.1 The Algorithm

We propose a new method named Conditional Angle Partition Tree (CAPT) for analyzing MD data, see Figure 1 for an overview. Intuitively, CAPT groups conformations into metastable states by recursively using the information from dihedral angles of the backbone chain with high partition scores. A higher partition score implies a higher energy barrier. The output of CAPT is a hierarchical partition tree of the conformations in the dihedral angle space. The boundaries between different partitions provide the locations of free energy barriers. CAPT also reports stable structures, which correspond to local minima of free energy basins. Details on CAPT are referred to **Methods**.

Comparing with previous methods, CAPT has following advantages: Firstly, CAPT operates directly on conformations instead of on microstates, thus we avoid the computing burden and inaccuracy caused by K-means/K-medoids in the splitting step. This advantage is only over the two-step clustering methods. Secondly, by incorporating the transition probability (dynamical similarity) into tree construction, CAPT considers simultaneously in each iteration the dynamical similarity and local geometrical similarity. With this feature, we believe that similarity in kinetic (dynamic) distances between different conformations is obtained from partial similarity in geometric distances. Thirdly, CAPT recognizes the number of metastable states automatically with the aid of local density, which is proportional to the free energy of the metastable state. However, many of the previous methods take this unknown number as input for analysis. Fourthly, CAPT actually performs divisive hierarchical clustering on all conformations using both dynamic and structural information, and thus can run in parallel, which is different from methods based on agglomerative clustering. Specifically, the branches of the partition tree can be constructed in parallel. For example, we can set a relatively higher value for *P*_*c*_, say 0.9, and get a tree. Next, we run CAPT in parallel on each end node of the tree with a smaller *P*_*c*_, say 0.5, and combine the subtrees to get the final partition tree. This will save computational time of CAPT. Fifthly, Combining with results from local density analysis, CAPT gives an informative representation of the free energy landscape with a desired solution.

The idea of using torsion angles to represent a biomolecules, i.e., assuming that the backbone chain structure is the main factor for forming major energy wells, has been used in some previous works, see for example, [52], [53]. Although the assumption behind CAPT that the energy wells are determined by backbone of the protein is supported by recent results [36], [37], [39], [43], [44], [52], it still may fail in some cases [54], [55]. In such cases, methods such as the optimal folding coordinate [56] and the transition disconnectivity graph [19], [20] may be tried.

### 3.2 Application to Alanine dipeptide

To explore the advantages of CAPT through its comparison with PCCA, PCCA+, MPP and Gibbs sampling method, we first apply CAPT to a benchmark dataset, the MD dataset of Alanine dipeptide. Details of analyzing Alanine dipeptide are presented for understanding the way that CAPT works. Alanine dipeptide is a simple and well-understood structure, whose molecular structure is given in Figure 2. The MD trajectory data of Alanine dipeptide is taken from [57], which consists of 974 20-ps NVE simulations with conformations stored every 0.1 ps. Thus there are 194800 conformations in the dataset. The detailed simulation information can be found in [57].

**Fig. 2.**
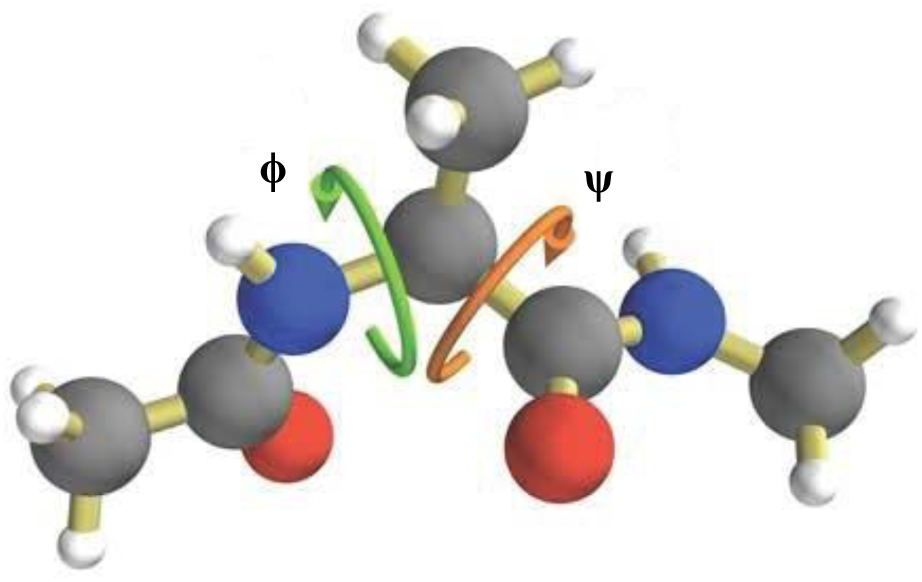
Structure of Alanine dipeptide. In the figure, the grey, red and blue atoms represent carbon, oxygen and nitrogen, respectively.

#### 3.2.1 Settings for Competitive Methods

Methods belonging to the two-step procedure, such as PCCA^∗^, PCCA+^*†*^, MPP^*‡*^, TRDG, MVCA and Gibbs sampling, have the same splitting step but different lumping steps. To help them eliminate the inaccuracy caused by their original splitting method on their final results, we used a grid clustering method on the angle space (See Supplementary Section 3.2) to cluster the conformations into microstates. Note that the grid clustering method provides much better clustering results than K-mean algorithm, thus favoring these two-step methods in comparison with our method. Here, we do not compare CAPT with APM since it is similar to PCCA/PCCA+ with the grid clustering method in the splitting step. Detailed settings for these four methods are given in Supplementary Section 3.2.

#### 3.2.2 CAPT Outperforms Competitive Methods

Alanine dipeptide has two torsion angles, *ϕ* and *ψ*. Thus, we can represent each conformation by these two angles, and it has 6 stable states with ideal landscape presented in Figure 4A. The histograms of *ϕ* and *ψ* are given in Figure 1(II) and the distribution of all conformations in (*ϕ, ψ*) space is given in Supplementary Figure 3. We run CAPT on the Alanine dipeptide data with *S*_*c*_ = 0, *S*_0_ = 500, *P*_*c*_ = *P*_0_ = 0.6 and the Gaussian kernel. A sensitivity analysis for these settings is given in Supplementary Section 3.5. The resulting partition tree is shown in Figure 3, which gives a estimator of partitions as (*B*_1_,, *B*_6_) = (*S*023, *S*022, *S*021, *S*013, *S*012, *S*011) (Transition matrix between them is shown in Supplementary Table 2). Thus, we conclude that Alanine dipeptide has 6 metastable states, which is consistent with the ground truth.

**Fig. 3.**
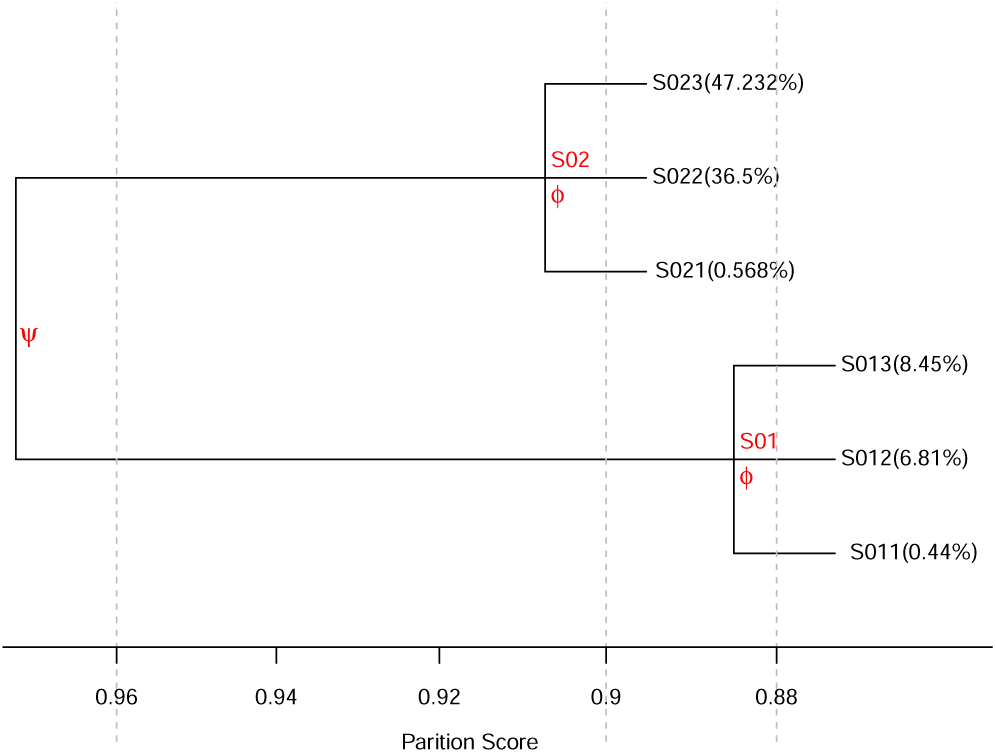
Partition tree of Alanine dipeptide. CAPT first uses *ψ* to partition all conformations into two child clusters, S01 and S02. For both S01 and S02, CAPT further uses *ϕ* to partition them each into three child clusters, as shown in the tree.

**Fig. 4.**
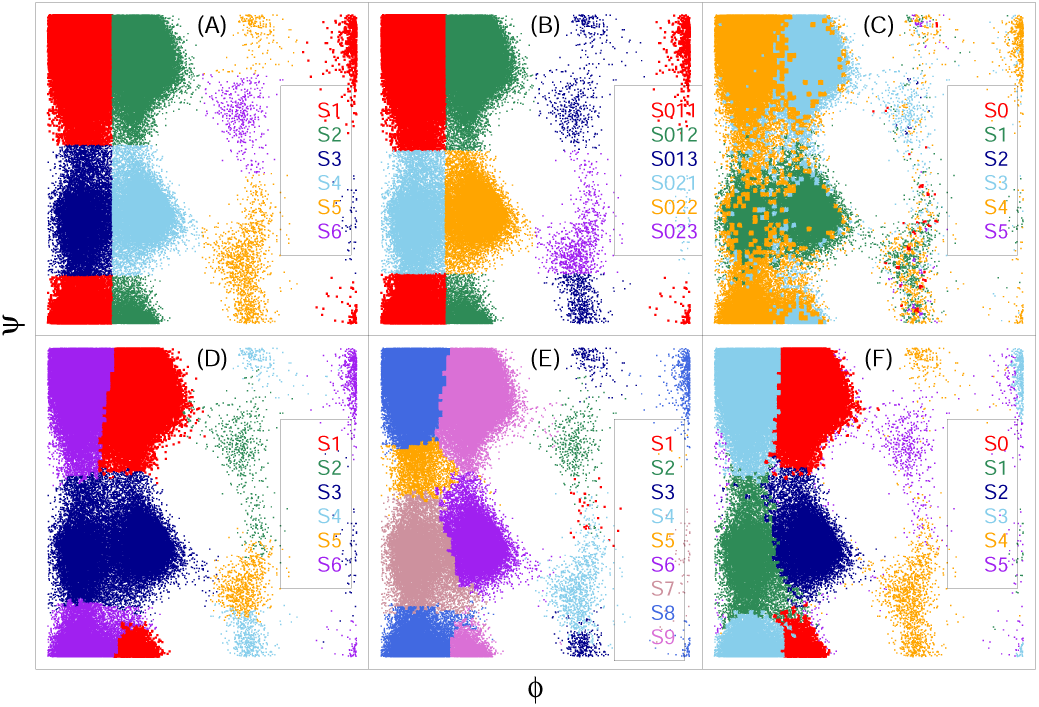
Free energy landscape of Alanine dipeptide obtained by different methods. Different colors in each sub-figure denote different clusters. (A) Benchmark results [57]; (B) CAPT; (C) MPP; (D) PCCA; (E) PCCA+; (F) Gibbs.

The free energy landscapes of Alanine dipeptide obtained by different methods are given in Figure 4. The corresponding transition matrix of each clustering is given in Supplementary Table 1-6. The free energy landscape plots in Figure 4 show that the result of CAPT is the one most similar to the benchmark result [57]. For a numerical comparison of results from different methods with the benchmark, we use the Adjusted Rand Index (ARI) [59] to quantify the similarity between two clusterings. If two clustering labels (*C*_1_, *C*_2_) are equivalent, we have ARI(*C*_1_, *C*_2_) = 1. If they are totally different, ARI(*C*_1_, *C*_2_) = 0. ARIs between cluster labels obtained by CAPT, PCCA, PCCA+, MPP, Gibbs sampling method and the benchmark cluster labels are given in Table 1, which also shows the corresponding standard errors estimated by bootstrapping the MD trajectories for 100 times. Table 1 shows that the cluster labels obtained by CAPT are the closest to the benchmark clustering and CAPT produces statistically significantly better ARI than other methods. The obvious differences among the landscapes shown in Figure 4 also support this finding. But since the difference in ARI between CAPT and Gibbs is only 0.035, one may argue that the difference is not scientifically meaningful. Thus, we suggest to also compare the results from different aspects other than ARI. As a result, CAPT performed much better on Alanine dipeptide than PCCA, PCCA+, MPP and Gibbs sampling method from three points of view: ARI (see Table 1), the free energy landscape (see Figure 4) and the transition matrix (Supplementary Table 1-6). Comparing with results from TRDG (see Supplementary Section 3.6) and MVCA (SI Figure 8), we arrive at the same conclusion.

**TABLE 1.** ARI (with standard error in the bracket) between the benchmark clustering and the cluster label from each method.

| ARI | CAPT | MPP |
| --- | --- | --- |
| Benchmark | 0.987 (0.0005) | 0.723 (0.0045) |
| PCCA | PCCA+ | Gibbs |
| 0.906 (0.0027) | 0.923 (0.0017) | 0.952 (0.0011) |

In summary, CAPT shows its advantage over other methods by incorporating the dynamical similarity into tree construction that it is computational efficient and gives an better estimate on number of macrostates.

#### 3.2.3 Discussion on Results from Competitive Methods

If we simply evaluate the results by the sum or the mean of diagonals of the transition matrix, which is the objective function of PCCA, the results given by PCCA is the best (see Supplementary Table 1-6). However, the free energy landscapes shown in Figure 4 suggest that the objective function of PCCA is ineffective. All these together suggest that mathematical objective of PCCA and PCCA+ may not be biological meaningful.

The Gibbs sampling method lumping the microstates into macrostates based on Poisson model shows some advantages over PCCA, PCCA+ and MPP as shown in Figure 4. But when looking into the transition matrix (Supplementary Table 6) from the Gibbs sampling algorithm, we find that the self-transition probability of state S5 is 0.527, which is unreasonable and may be caused by ignoring the geometric similarity in lumping step.

The transition matrix between clusters obtained from MPP looks unreasonable. The mean and minimal of diagonals are as low as 0.4449 and 0.0241, respectively, which defies the energy well explanation of a cluster.

In addition, we can see from Figure 4(C) that some conformations belonging to S4 are clustered into S3 by MPP, since there is a most probable pathway that starts from them and goes through a conformation with the lowest free energy belonging to S3. This result suggests that the principle used by MPP in dynamical clustering may be problematic.

PCCA, PCCA+ and MPP can not get the reasonable clustering labels even when we set the number of clusters from it as the true number of clusters of Alanine dipeptide and a better splitting of conformations into microstates. Similar to PCCA and PCCA+, the disadvantage of Gibbs sampling method is its requirement of a pre-defined number of clusters. Here we set it as the true number of clusters, however, it is unknown to us in practice. For Alanine dipeptide, we also run PCCA+ and Gibbs sampling algorithm on different settings of number of clusters produced from them. The results show that these algorithms depend strongly on the pre-defined number of clusters. For details, we refer to Supplementary Section 3.5. TRDG is a graph clustering algorithm, where the energy barrier between microstates is estimated by the minimum-cut between them in the graph. The results in Supplementary Figure 7 show that TRDG can recognize a part of local stable states but microstates from different benchmark clusters are mixed up. This phenomenon is also observed in metric disconnectivity graphs [60]. Another important point for TRDG is that it is difficult to judge from its results (SI Figure 7B-7E) how many metastable states are there, whereas CAPT provides an accurate estimate. It is well known that, disconnectivity graph is predominantly used for visualisation purposes and is highly sensitive to the choice of the threshold, which may explain partially its non-ideal performance on the MD data of Alanine dipeptide.

### 3.3 Application to HP35 double mutant Nle/Nle

In this section, we apply CAPT to HP35 double mutant Nle/Nle, whose structure is given in Figure 5. Its free energy landscape has been reported in [44] based on MPP, in [61] by both conventional and replica exchange molecular dynamics simulations, and in [56], [62] by constructing the optimal reaction coordinate. The data we used here is from [63], a ≈ 300 *µ*s long trajectory at 380 K, which shows 140 folding and unfolding events. The data contains ≈ 1.5 × 10^6^ snapshots with a time step of 200 ps.

**Fig. 5.**
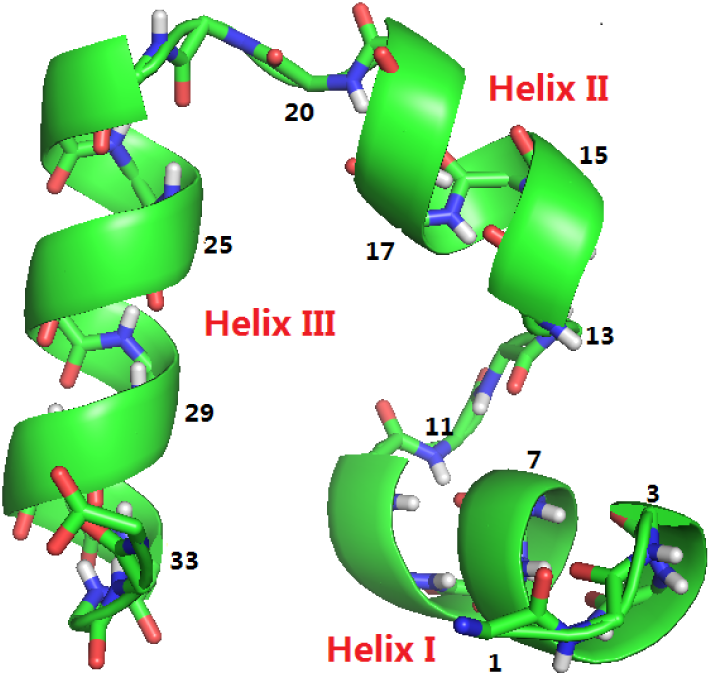
The structure of HP35 Nle/Nle determined by x-ray crystallography (PDB code 2F4K). The backbone is shown as cartoon. Oxygen is colored in red, C^*α*^ is colored in green with a number beside, and nitrogen is colored in blue.

This is a complex biomolecule, and CAPT may produce many small meaningless clusters. Local density is used to identify the stable structures and thus the number of marcrostates. The conformation with highest local density in the cluster is defined as the most stable one. We use two different local densities: (1) local density within the corresponding cluster (LDc) and (2) local density over all conformations (LDa). If the clusters are well separated, LDc is the same as LDa. Thus, a comparison between LDc and LDa tells the quality of the clustering.

#### 3.3.1 Results from CAPT

We run CAPT with *P*_*c*_ = *P*_0_ = 0.7, *S*_0_ = *S*_*c*_ = 10000 and Gaussian kernel. More discussion on selecting *S*_*c*_ and *S*_0_, and the sensitivity analysis on *P*_*c*_ are given in Supplementary Section 4.3. The full partition tree under this setting is presented in Supplementary Figure 15, and the corresponding results of LDc and LDa are given in Supplementary Table 19, where the threshold *d*_0_ for computing local density is set as 0.1757350, the 0.05% quantile of the one-step MAD (see **Methods** for definition, and details on selecting *d*_0_ is given in Supplementary Section 4.1). According to these results, we find 6 stable clusters, and the structures of their centers are given in Figure 7 with the transition matrix given in Supplementary Table 16. The structure S6, the most stable one, is exactly the native structure shown in Figure 5. We show the main partition tree between these 6 clusters in Figure 6, which tells the main difference between these 6 stable structures shown in Figure 7. More specifically, as shown in Figure 6, the energy barrier between {S1, S2} and {S3, S4, S5, S6} is caused by *ϕ*_2_; the energy barriers between S1 and S2, S3 and S4, S5 and S6, are all caused by *ϕ*_3_; and the energy barrier between {S3, S4} and {S5, S6} is caused by *ψ*_2_. This information is consistent with that contained in structures shown in Figure 7.

**Fig. 6.**
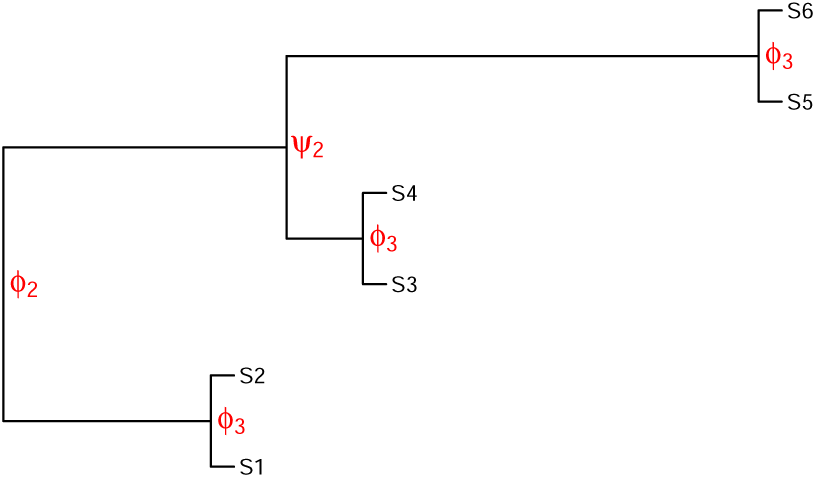
Relative free energy landscape between top 6 stable clusters with the key energy barrier.

**Fig. 7.**
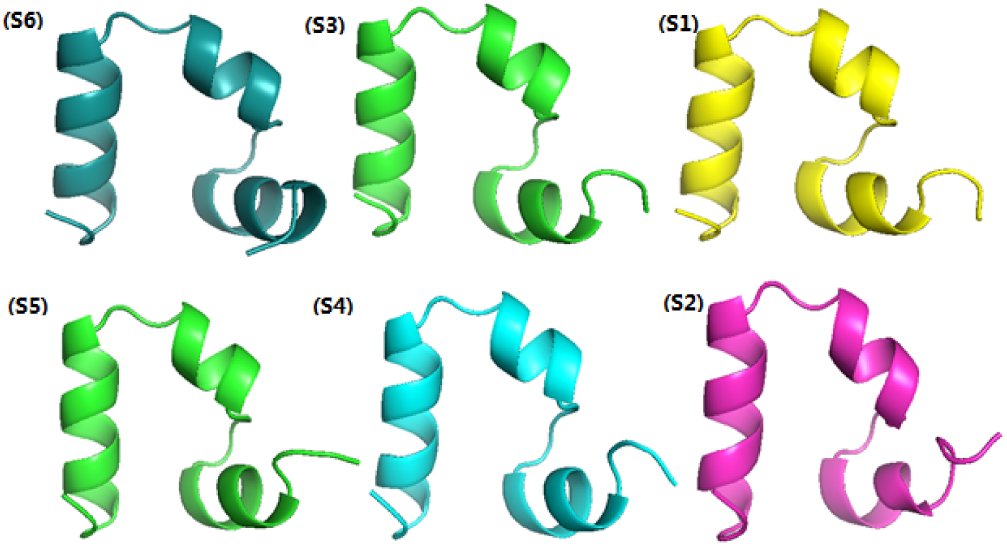
Top 6 most stable structures (with largest LDc with *d*_0_ = 0.1757350), sorted by their LDc in decreasing order from left to right and up to below. The labels are given according to the partition tree in Figure 6. Details are in Supplementary Material.

Since structures presented in Figure 7 are similar, we conclude that these are mainly native structures or intermediate structures. To fully understand the free energy landscape of HP35, we look inside into the full partition tree presented in Supplementary Figure 15 and the LDc/LDa in Supplementary Table 19, and get the refined partition tree as shown in Figure 8, which is a sub-tree of that in Supplementary Figure 15. We show in Figure 9 three examples of conformations in clusters L1-L5, L7 and L8. The transition matrix between these 9 clusters (metastable states) is shown in Table 2.

**Fig. 8.**
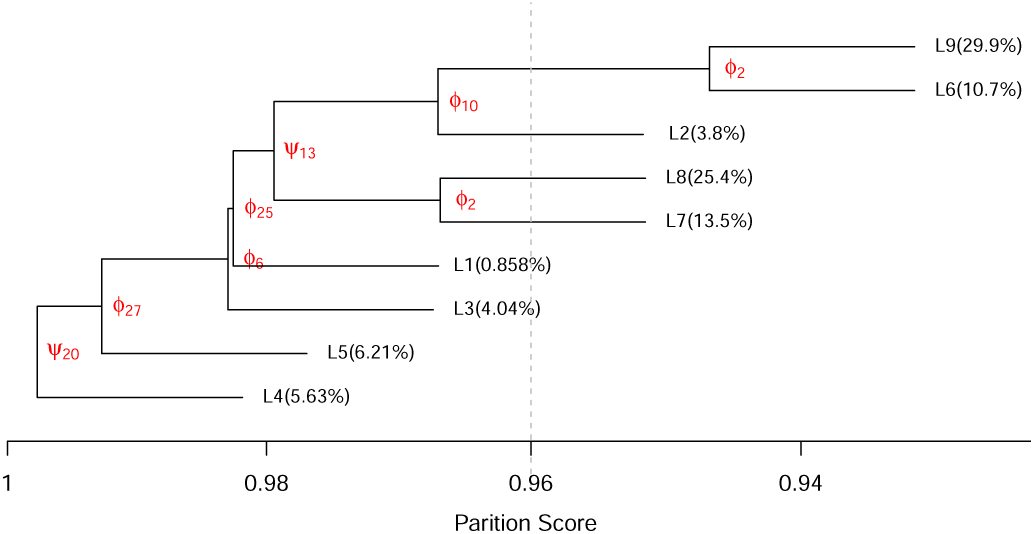
The final partition tree of HP35 Nle/Nle obtained with the aid of structures information in Figure 7 and LDc/LDa in Supplementary Table 19. The number in the parentheses is the population of the cluster. This partition tree is a sub-tree of that in Supplementary Figure 15. Comparing with the tree presented in Figure 6, it is clear that {S5, S6, S4, S3} belongs to cluster L9, {S1, S2} belongs to cluster L6.

**Fig. 9.**
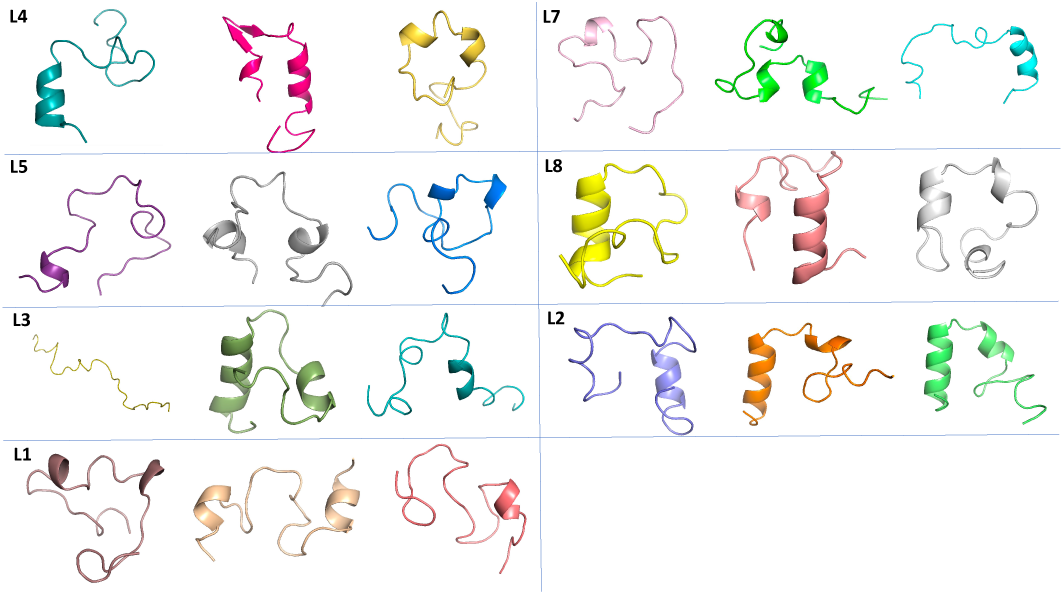
The examples of conformations in cluster L1-L5, L7 and L8. We present 3 examples for each cluster.

**TABLE 2.**
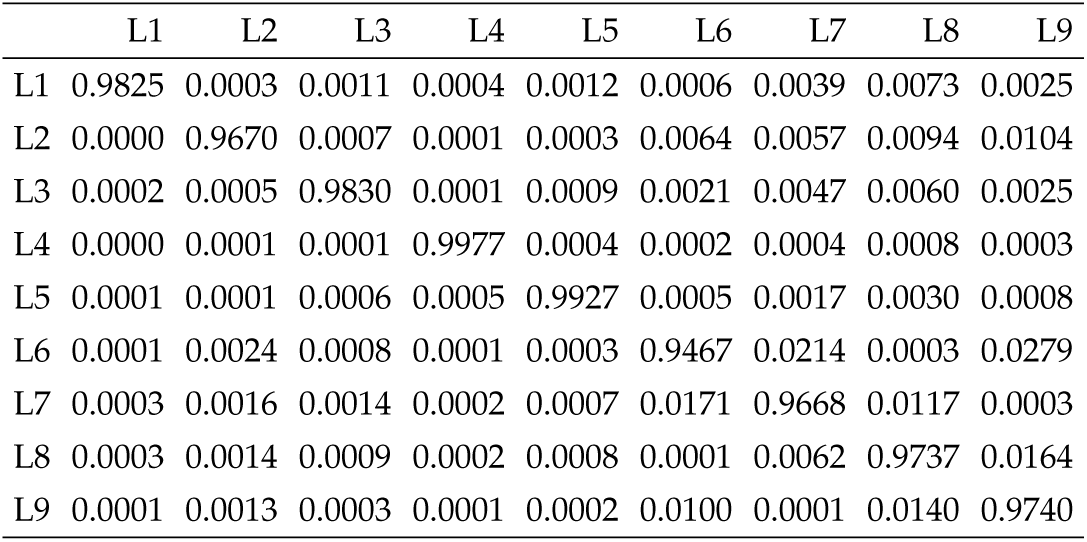
Transition matrix between clusters in Figure 8

|  | L1 | L2 | L3 | L4 | L5 | L6 | L7 | L8 | L9 |
| --- | --- | --- | --- | --- | --- | --- | --- | --- | --- |
| L1 | 0.9825 | 0.0003 | 0.0011 | 0.0004 | 0.0012 | 0.0006 | 0.0039 | 0.0073 | 0.0025 |
| L2 | 0.0000 | 0.9670 | 0.0007 | 0.0001 | 0.0003 | 0.0064 | 0.0057 | 0.0094 | 0.0104 |
| L3 | 0.0002 | 0.0005 | 0.9830 | 0.0001 | 0.0009 | 0.0021 | 0.0047 | 0.0060 | 0.0025 |
| L4 | 0.0000 | 0.0001 | 0.0001 | 0.9977 | 0.0004 | 0.0002 | 0.0004 | 0.0008 | 0.0003 |
| L5 | 0.0001 | 0.0001 | 0.0006 | 0.0005 | 0.9927 | 0.0005 | 0.0017 | 0.0030 | 0.0008 |
| L6 | 0.0001 | 0.0024 | 0.0008 | 0.0001 | 0.0003 | 0.9467 | 0.0214 | 0.0003 | 0.0279 |
| L7 | 0.0003 | 0.0016 | 0.0014 | 0.0002 | 0.0007 | 0.0171 | 0.9668 | 0.0117 | 0.0003 |
| L8 | 0.0003 | 0.0014 | 0.0009 | 0.0002 | 0.0008 | 0.0001 | 0.0062 | 0.9737 | 0.0164 |
| L9 | 0.0001 | 0.0013 | 0.0003 | 0.0001 | 0.0002 | 0.0100 | 0.0001 | 0.0140 | 0.9740 |

Comparing with the tree presented in Figure 6, we see that S5, S6, S4, S3 belongs to cluster L9, {S1, S2} belongs to cluster L6. According to structures shown in Figure 7, state L9 can be defined as the native state, and state L6 is the intermediate state. In addition, according to Figure 8, whether an unfolded conformation belonging to L4 can evolved into a conformation in native state L9 is determined by torsion angles {*ϕ*_2_, *ϕ*_6_, *ϕ*_10_} in Helix I, {*ψ*_13_} in Helix II, {*ϕ*_25_, *ϕ*_27_} in Helix III, and *ψ*_20_ in the middle of the structure in Figure 5. Combining results in Figure 9 and Figure 8, we see that the folding of a molecule starts from the easier part having a lower energy barrier to the harder part with a higher energy barrier. Furthermore, results presented in Figure 9 and Table 2 validate the statement that geometric similarity may not imply kinetic (dynamic) similarity, see for example, [52].

#### 3.3.2 Comparing with results from MPP

In this section, we compare the results from CAPT with that from MPP, where the clustering results from MPP is obtained from the authors of [44]. Table 3 shows the maximal LDc of clusters found by MPP and CAPT (more details are given in Supplementary Table 14), where U is the unfolded state, I1 and I2 are the intermediate states, and N1 and N2 are native states. From these results, we see that the maximal LDc of cluster U and N2 found by MPP is less than 100, which is smaller than the minimum maximal LDc of clusters found by CAPT. This indicates that the result obtained by MPP is somehow unreasonable. In addition, as shown in Table 3 and Supplementary Table 14, the LDc and LDa of the most stable structures in cluster I1 and I2 are not equal even when the threshold *d*_0_ is as small as the 0.005% quantile of one-step MAD. This further indicates the drawback of MPP.

**TABLE 3.** LDc and LDa of stable conformations found by MPP and CAPT when *d*_0_ = 0.1757350

| MPP [44] |  |  | CAPT |  |  |
| --- | --- | --- | --- | --- | --- |
| Cluster | LDC | LDa | Cluster | LDC | LDa |
| I1 | 6766 | 6854 | S6 | 6854 | 6854 |
| N1 | 3789 | 3790 | S3 | 3662 | 3662 |
| U | 67 | 67 | S1 | 3167 | 3167 |
| I2 | 816 | 839 | S5 | 1321 | 1321 |
| N2 | 8 | 8 | S4 | 834 | 834 |
| - | - | - | S2 | 763 | 763 |

Furthermore, we show in Table 4 the relationship between the clusters from MPP and CAPT. According to the results, members in clusters I1, N1, I2 and N2 mainly belong to clusters L6 and L9, which are native (or intermediate) metastable states. This situation is similar to that occurred in Alanine dipeptide (see Figure 4): one metastable states may be divided by MPP into several states (clusters). Interestingly, members of clusters L1 - L5, L7 and L8, with about 59.4% population, mainly belong to the cluster U, the unfolded state. This is consistent with the partition tree shown in Figure 8, which is a representation of free energy landscape. According to the partition tree, we may understand that the unfolded state can be divided into 7 unfolded sub-states. Also, one can divide, according to the full partition tree in SI Figure 16, the native cluster L9 further to see detailed free energy landscape for the native state. This is consistent with the fact that the free energy landscape is rugged. The members in cluster U are separated in clusters L1-L9, which explains why the most stable structure in U found by local density is different from that in [44]. Importantly, many conformations belong to the native state are recognized mistakenly by MPP as unfolded states (see Table 3 and 4, Figure 6 and 8).

**TABLE 4.** Number of conformations in intersection between clusters from MPP and CAPT

|  | L1 | L2 | L3 | L4 | L5 | L6 | L7 | L8 | L9 |
| --- | --- | --- | --- | --- | --- | --- | --- | --- | --- |
| I1 | 3 | 114 | 576 | 18 | 33 | 5911 | 11 | 40 | 115907 |
| N1 | 0 | 30 | 190 | 0 | 5 | 29142 | 4 | 9 | 84056 |
| U | 12828 | 57139 | 57772 | 84739 | 92738 | 87625 | 203529 | 382920 | 172981 |
| I2 | 3 | 29 | 2412 | 1 | 5 | 31336 | 10 | 13 | 56452 |
| N2 | 120 | 40 | 8 | 241 | 888 | 7649 | 1 | 4 | 21860 |

In summary, CAPT gives an informative representation of the free energy landscape, and the key difference between each cluster in the partition tree is well understood. Importantly, with the aid of local density analysis, the free energy landscape obtained from CAPT can explore the landscape with a required resolution at the part of interest. For example, we can just take clusters L1-L5, L7 and L8 as one unfolded state or treat them as different unfolded states. Also, we may take S6 as the native state, and treated S1-S5 as intermediate states or just take L9 as the native state and L6 as the intermediate state.

## 4 Conclusion

We introduced a new method for deciphering the free energy landscape of the conformational space of biomolecules. Our method makes use of both the three dimensional structure of each conformation and the dynamic information between conformations to group conformations into metastable states. A special characteristics of our new method is its usage of the local structure information, more specifically the discriminant angles, to construct a classification tree. The resulting clusters have both high local densities and low between-group moves, which fits the concept of energy wells. When compared with other methods, our method achieved better results as judged by the benchmark or the local density of stable structures.

CAPT aims to learn the metastable states of biomolecules (proteins) from MD data. Metastable states are defined as states with a self-transition probability larger than a given threshold *P*_*c*_ ∈ [0.5, 1]. Compared with popular two-step procedures such as PCCA/PCCA+, MPP and Gibbs sampling method, CAPT works on a different framework. Specifically, those two-step methods take the problem of exploring the free energy landscape as a clustering problem (clustering conformations into macrostates by two steps), while CAPT takes it more like a non-supervised classification problem. The advantage of this new view is that it avoids the local optima of K-means algorithm that may result in bad microstates for complex biomolecules, and the tedious calculation of K-means. Importantly, CAPT has many other advantages over the other methods. Molecular dynamical simulations is an important technique for understanding big biomolecules but it is time-consuming. Sometimes, other methods like discrete path sampling [64] are used for sampling the trajectories, where CAPT is still applicable.

For complex molecules or proteins such as HP35, there is no ground truth to compare with. Researchers tend to validate the result in terms of self-consistency. For example, we validate our results and compare them with those in [44] by using the local density within the cluster (LD_*c*_) and local density over all conformations (LD_*a*_) of the metastable states (see Table 3), and using the simple principle that a metastable state has a high local density. A crosscheck of the outputs from other related energy landscape studies might be possible, but challenging to be objective in the absent of ground truth. Also, benchmark datasets are in great need for understanding and comparing the performance of different methods, which is an important research topic for this area.

Currently our method is an intuitive data-driven approach. One way to improve over our method is to introduce a generative probabilistic model to describe the free energy landscape more principally, which needs detailed modelling of the global similarity and the local similarity among conformations. The energy barrier casts a non-continuous factor to the model, which will be the major challenge for the probabilistic modeling. Furthermore, the detail quantification of free energy minima or barrier might be pursued but not dealt in the current paper. These interesting topics are left as the future work.

## Data availability

Trajectory data of HP35 Nle/Nle can be obtained from https://www.deshawresearch.com/downloads/ download trajectory pnas2012.cgi/ with help from authors of [63].

## Code availability

The code for CAPT algorithm is available at https://jiangdata.github.io/resources/capt.zip.

## Acknowledgements

We would like to thank Abhinav Jain and Gerhard Stock for kindly sharing their detailed results in [44] with us, and thank Prof. David J. Wales from Department of Chemistry at University of Cambridge for his help on drawing the disconnectivity graph. This work is partially supported by one grant from the National Science Foundation of USA (DMS-1811920), three grants from the Research Grants Council of the Hong Kong SAR, China (400913, 14203915, 14173817), and one grant from The National Natural Science Foundation of China (11901517).

## Footnotes

∗. The code is from MSMbuilder 2.0 [58]

†. The code is provided by Marcus Weber, author of [17]

‡. The code is provided by its authors

## Notes

### Competing Interest Statement

The authors have declared no competing interest.

### Summary of Updates

Authorship updated and supplemental files updated.

