## supplemental Material for "Revealing Free Energy Landscape from MD Data via Conditional Angle Partition Tree"

#### 1 Overview of MD data and transition matrix

Figure 1 shows an overview of MD trajectory data and related transition matrix in two-step clustering procedure.

#### 2 Data Preparation

The input of CAPT is a set of MD trajectories. We use the R package 'bio3d' [1] to get all torsion angles of all frames of the MD trajectories. In this paper, we only use the backbone torsion angles, i.e., $\phi's$ and $\psi's$.

#### 3 Details about analyzing Alanine dipeptide

##### 3.1 Visualization of Alanine dipeptide data

Figure 2 shows the histograms of angles $(\psi, \phi)$ of Alanine dipeptide. Figure 3 shows the distribution of the frames of Alanine dipeptide data in the angle phase.

##### 3.2 Settings for PCCA, PCCA+, MPP and Gibbs

When using PCCA/PCCA+, one should specify the range of the number of clusters, i.e., the maximal number of clusters $n_{max}$ and minimal number of clusters $n_{min}$. We set $n_{max} = n_{min} = 6$ for PCCA, and $n_{max} = 9$ and $n_{min} = 3$ for PCCA+. The Gibbs sampling algorithm is also run with the true number of clusters.

To use PCCA, PCCA+, MPP and Gibbs sampling method, one has to cluster frames into microstates first. In the simple case of Alanine dipeptide which only has two torsion angles, we use a grid method in the angle space to group frames to microstates. More specifically, we partition the whole space $[-\pi, \pi] \times [-\pi, \pi]$ into a $80 \times 80$ grid, and take each small grid as a microstate. The center of each small grid cell is treated as the center of this microstate. In fact, by dividing the $(\phi, \psi)$ space into 6400 bins, we get many empty bins and remove them from our analysis. We take each non-empty bin as a microstate, and map each conformation in the trajectory to the corresponding microstate, then we get trajectories of microstates, and the transition matrix between microstates.

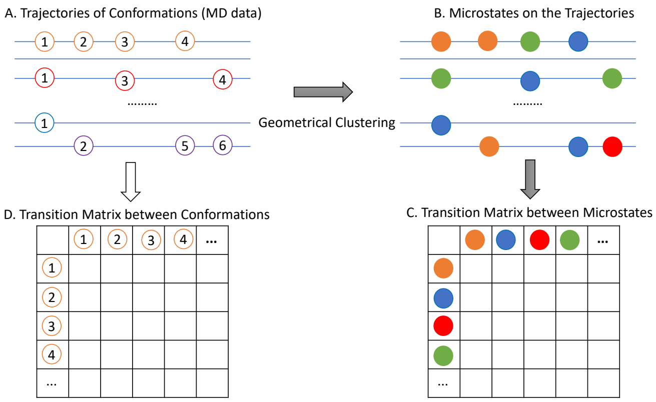


Fig. 1. MD trajectory data and related transition matrix. (A) shows the trajectories of conformations (denoted by empty circles) obtained from molecular dynamics simulation. Note that circles in different color denote different conformations in different trajectories. (B) shows the trajectories of microstates (denoted by solid colored circles) by mapping conformations to microstates through geometrical clustering. (C) shows the transition matrix, $T(m \times m)$, between microstates obtained from trajectories of conformations in sub-figure (B), where $m$, usually no more than thousands, is the number of microstates obtained by geometrical clustering in the splitting step. The entry of this matrix, $T_{ij}$, is the number of jumping from microstate $i$ to microstate $j$ along the trajectories. That is, $T_{ij}$ is the number of times we observe in all trajectories that the microstate $i$ is followed by microstate $j$. (D) shows the transition matrix, $K(n \times n)$, between conformations obtained from trajectories of conformations in sub-figure (A), where $n$, usually at the magnitude of millions, is the number of conformations observed in MD simulations. The entry of this matrix, $K_{ij}$, is the counts of jumping from conformation $i$ to conformation $j$ along the trajectories. That is, $K_{ij}$ is the number of times we observe in all trajectories that the conformation $i$ is followed by conformation $j$. Since any two conformations along the trajectories are different, $L(s-1)$ of the entries of $K$ are 1, and others are 0, thus non-informative for detecting stable structures, where $L$ is the number of trajectories, $s$ is the number of conformations in each trajectory, and $n = L \times s$.


- *Hangjin Jiang is with the Center for Data Science, Zhejiang University.*
- *Han Li is with College of Economics, Shenzhen University.*
- *Wing Hung Wong is with the Department of Statistics, Stanford University.*
  **
- *Xiaodan Fan is with the Department of Statistics, The Chinese University of Hong Kong*
  **

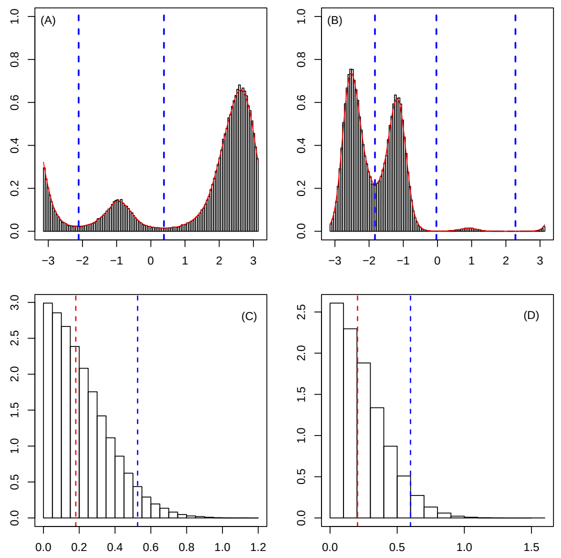


Fig. 2. Histograms of angles ($\psi, \phi$) of Alanine dipeptide. (A) and (B) are the histograms of $\psi$ and $\phi$, respectively. (C) and (D) are the histograms of the one-step distance of $\psi$ and $\phi$, respectively. In (A) and (B), the blue dashed lines show the partition positions found by CAPT. In (C) and (D), the blue and red dashed lines correspond to the 95% and 50% quantiles of the distribution, respectively.

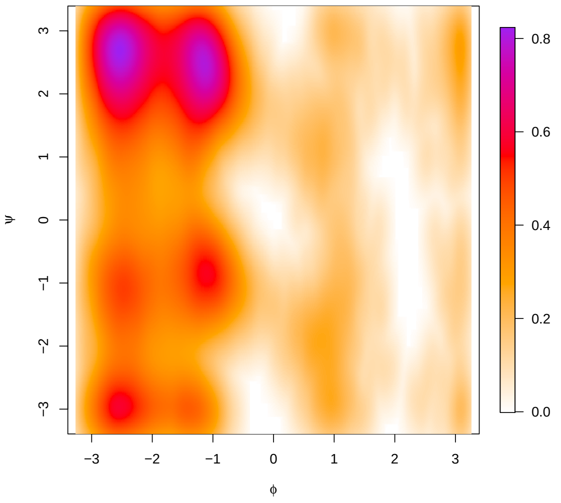


Fig. 3. Scatter plot of frames of Alanine dipeptide data in the angle phase. The Alanine dipeptide is a small biomolecule with only two torsion angles, say, $\phi$ and $\psi$, and its free energy landscape is fully determined by these two angles.

##### 3.3 Transition matrix from different methods

Table 1 shows the transition matrix of the benchmark clusters of Alanine dipeptide, Table 2 shows results from CAPT, Table 4 shows results from PCCA, Table 3 shows that of clusters of Alanine dipeptide obtained by MPP, Table 5 show results from PCCA+, Table 6 show results from Gibbs sampling method.

TABLE 1
Transition matrix of the benchmark clusters of Alanine dipeptide

| | S1 | S2 | S3 | S4 | S5 | S6 |
|---|---|---|---|---|---|---|
| S1 | 0.9457 | 0.0477 | 0.0062 | 0.0004 | 0.0000 | 0.0000 |
| S2 | 0.0609 | 0.9365 | 0.0004 | 0.0021 | 0.0000 | 0.0002 |
| S3 | 0.0403 | 0.0021 | 0.8939 | 0.0636 | 0.0000 | 0.0000 |
| S4 | 0.0020 | 0.0090 | 0.0526 | 0.9356 | 0.0008 | 0.0000 |
| S5 | 0.0013 | 0.0013 | 0.0000 | 0.0098 | 0.9718 | 0.0158 |
| S6 | 0.0000 | 0.0401 | 0.0000 | 0.0000 | 0.0519 | 0.9080 |
| Sum of diagonals: 5.591479 | | | | | | |
| Mean of diagonals: 0.9319131 | | | | | | |
| Minimal of diagonals: 0.8939 | | | | | | |

TABLE 2
Transition matrix between clusters of Alanine dipeptide obtained by CAPT

| | S011 | S012 | S013 | S021 | S022 | S023 |
|---|---|---|---|---|---|---|
| S011 | 0.9461 | 0.0475 | 0.0000 | 0.0061 | 0.0003 | 0.0000 |
| S012 | 0.0613 | 0.9360 | 0.0002 | 0.0004 | 0.0021 | 0.0000 |
| S013 | 0.0027 | 0.0163 | 0.9074 | 0.0000 | 0.0000 | 0.0736 |
| S021 | 0.0407 | 0.0019 | 0.0000 | 0.8932 | 0.0642 | 0.0000 |
| S022 | 0.0018 | 0.0095 | 0.0000 | 0.0515 | 0.9364 | 0.0009 |
| S023 | 0.0000 | 0.0000 | 0.0975 | 0.0000 | 0.0176 | 0.8848 |
| Sum of diagonals: 5.50389 | | | | | | |
| Mean of diagonals: 0.917315 | | | | | | |
| Minimal of diagonals: 0.8848 | | | | | | |

##### 3.4 Discussion

If we compare the energy landscape of benchmark clusters of Alanine dipeptide with that obtained by CAPT, although ARI is as high as 0.987002, we can still find a small difference: the boundary between S013 and S023 in Figure 4(B) is a bit higher than that between S5 and S6 in Figure 4(A). This small difference is due to the order of angles used for partitioning. Table 7 gives the transition matrix when using different angles in the first step of CAPT for Alanine dipeptide. The $\psi$ column is the transition matrix between clusters obtained by partitioning all frames into 2 clusters according to the distribution of $\psi$, where the partition score is 0.9728. The $\phi$ column is the transition matrix between clusters obtained by partitioning all frames into 3 clusters according to the distribution of $\phi$, where the partition score is 0.9395. Thus CAPT prefers using $\psi$ to partition in the first step. That means that CAPT is a bit greedy in transition probability, which leads to the small discrepancy from the benchmark.

Alternatively, if CAPT had used $\phi$ first in constructing the partition tree, we would get the energy landscape as shown in Figure 4, which would be closer to the benchmark in Figure 4(A) (ARI=0.9873). The corresponding transition matrix is given in Table 8. This implies that there is room to improve the performance of CAPT if we can find a more suitable partition score.

##### 3.5 Sensitivity Analysis

As we can see from the results of PCCA+ in Figure 4, PCCA+ overestimates the number of clusters under the setting in Section 2.2 of this Supplementary Material. We try another setting for PCCA+, $n_{max} = 5$ and $n_{min} = 7$, denoted as PCCA+′. We also run the Gibbs sampling algorithm with 5 and 7 clusters, denoted by Gibbs′ and

TABLE 3
Transition matrix between clusters of Alanine dipeptide obtained by MPP

| | S0 | S1 | S2 | S3 | S4 | S5 |
|---|---|---|---|---|---|---|
| S0 | 0.0526 | 0.3289 | 0.0000 | 0.1447 | 0.4342 | 0.0395 |
| S1 | 0.0013 | 0.8124 | 0.0001 | 0.0460 | 0.1391 | 0.0011 |
| S2 | 0.0000 | 0.2778 | 0.0556 | 0.6667 | 0.0000 | 0.0000 |
| S3 | 0.0001 | 0.0159 | 0.0002 | 0.8503 | 0.1335 | 0.0000 |
| S4 | 0.0003 | 0.0330 | 0.0000 | 0.0918 | 0.8745 | 0.0005 |
| S5 | 0.0241 | 0.3012 | 0.0000 | 0.0361 | 0.6145 | 0.0241 |

Sum of diagonals: 2.669393
Mean of diagonals: 0.4448989
Minimal of diagonals: 0.0241

TABLE 4
Transition matrix between clusters of Alanine dipeptide obtained by PCCA

| | S1 | S2 | S3 | S4 | S5 | S6 |
|---|---|---|---|---|---|---|
| S1 | 0.9352 | 0.0003 | 0.0018 | 0.0000 | 0.0000 | 0.0626 |
| S2 | 0.0477 | 0.9131 | 0.0000 | 0.0068 | 0.0324 | 0.0000 |
| S3 | 0.0042 | 0.0000 | 0.9752 | 0.0000 | 0.0004 | 0.0202 |
| S4 | 0.0000 | 0.0032 | 0.0000 | 0.9104 | 0.0816 | 0.0048 |
| S5 | 0.0000 | 0.0269 | 0.0175 | 0.0672 | 0.8884 | 0.0000 |
| S6 | 0.0508 | 0.0000 | 0.0068 | 0.0000 | 0.0000 | 0.9424 |

Sum of diagonals: 5.564797
Mean of diagonals: 0.9274662
Minimal of diagonals: 0.8884

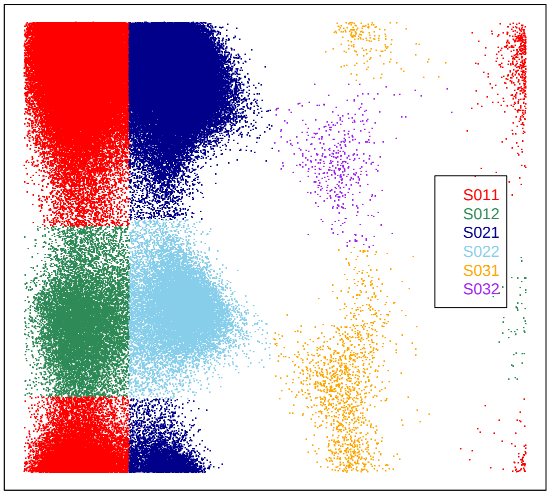


Fig. 4. Energy landscape of Alanine dipeptide obtained from CAPT by using firstly $\phi$ to partition the angle space.

Gibbs$''$. Transition matrice from PCCA+$'$, Gibbs$'$ and Gibbs$''$ are shown in Table 9, Table 10 and Table 11. The clustering results are shown in Figure 5. All these results show that both PCCA+ and Gibbs sampling algorithm depend strongly on the pre-selected number of clusters. However, the number of stable states (clusters) is usually unknown to us and it is the key quantity to be estimates from the data.

As reported in [2], [3], [4], Dihedral PCA gives clear energy landscape of biomolecules. However, while we try it on Alanine dipeptide, its partitioning is not clear as shown in Figure 6.

For sensitivity analysis of our algorithm, we check the effect of parameters $P_0$ and $S_0$ as well as the kernel

TABLE 5
Transition matrix between clusters of Alanine dipeptide obtained by PCCA+

| | S1 | S2 | S3 | S4 | S5 | S6 | S7 | S8 | S9 |
|---|---|---|---|---|---|---|---|---|---|
| S1 | 0.7667 | 0.1333 | 0.0000 | 0.1000 | 0.0000 | 0.0000 | 0.0000 | 0.0000 | 0.0000 |
| S2 | 0.0096 | 0.9349 | 0.0096 | 0.0145 | 0.0000 | 0.0000 | 0.0000 | 0.0000 | 0.0313 |
| S3 | 0.0000 | 0.0045 | 0.8649 | 0.1194 | 0.0000 | 0.0000 | 0.0000 | 0.0068 | 0.0045 |
| S4 | 0.0030 | 0.0089 | 0.0523 | 0.9211 | 0.0000 | 0.0148 | 0.0000 | 0.0000 | 0.0000 |
| S5 | 0.0000 | 0.0000 | 0.0000 | 0.0000 | 0.6736 | 0.0412 | 0.0659 | 0.1643 | 0.0550 |
| S6 | 0.0000 | 0.0000 | 0.0000 | 0.0009 | 0.0069 | 0.9344 | 0.0539 | 0.0003 | 0.0037 |
| S7 | 0.0000 | 0.0000 | 0.0000 | 0.0000 | 0.0144 | 0.0596 | 0.8945 | 0.0296 | 0.0019 |
| S8 | 0.0000 | 0.0000 | 0.0000 | 0.0000 | 0.0052 | 0.0000 | 0.0046 | 0.9419 | 0.0483 |
| S9 | 0.0000 | 0.0001 | 0.0000 | 0.0000 | 0.0022 | 0.0009 | 0.0005 | 0.0601 | 0.9361 |

Sum of diagonals: 7.86813
Mean of diagonals: 0.8742366
Minimal of diagonals: 0.6736

TABLE 6
Transition matrix between clusters of Alanine dipeptide obtained by Gibbs Sampling method

| | S0 | S1 | S2 | S3 | S4 | S5 |
|---|---|---|---|---|---|---|
| S0 | 0.9349 | 0.0012 | 0.0018 | 0.0605 | 0.0001 | 0.0015 |
| S1 | 0.0058 | 0.8783 | 0.0666 | 0.0443 | 0.0002 | 0.0047 |
| S2 | 0.0073 | 0.0562 | 0.9313 | 0.0029 | 0.0010 | 0.0013 |
| S3 | 0.0477 | 0.0069 | 0.0007 | 0.9429 | 0.0000 | 0.0018 |
| S4 | 0.0033 | 0.0020 | 0.0113 | 0.0020 | 0.9515 | 0.0299 |
| S5 | 0.1374 | 0.0640 | 0.0284 | 0.1919 | 0.0509 | 0.5273 |

Sum of diagonals: 5.1662
Mean of diagonals: 0.8610
Minimal of diagonals: 0.5273

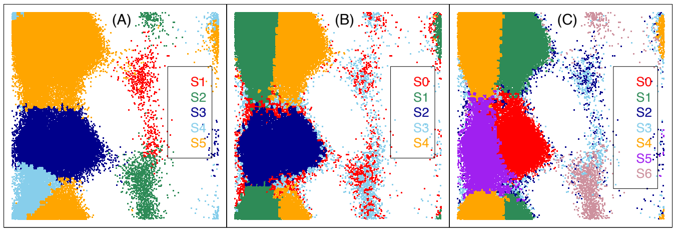


Fig. 5. Clustering result of Alanine dipeptide from PCCA+$'$ (A), Gibbs$'$ (B) and Gibbs$''$ (C).

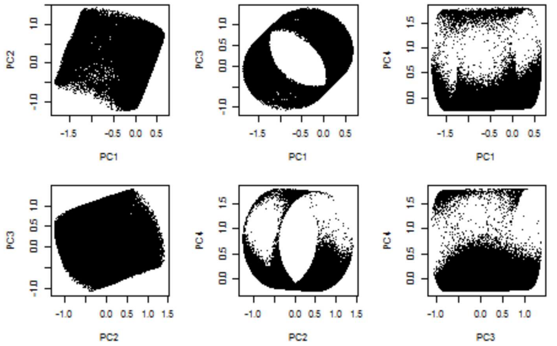


Fig. 6. The effect of Dihedral PCA on Alanine dipeptide. We apply Dihedral PCA to the Alanine dipeptide data, but do not see any pattern explored by PCA.

function while applying CAPT on Alanine dipeptide MD data. For the kernel function, we tried different kernel functions, such as Gaussian, von Mises and Epanechnikov,

TABLE 7
Transition matrix when using different partition schemes in the first step

| | $\psi$ | | | $\phi$ | | |
|---|---|---|---|---|---|---|
| | S1 | S2 | | S1 | S2 | S3 |
| S1 | 0.9949 | 0.0051 | S1 | 0.9496 | 0.0503 | 0.0000 |
| S2 | 0.0272 | 0.9728 | S2 | 0.0602 | 0.9395 | 0.0003 |
| - | - | - | S3 | 0.0015 | 0.0174 | 0.9811 |

TABLE 8
Transition matrix between clusters when first using $\phi$ to partition in CAPT

| | S011 | S012 | S021 | S022 | S031 | S032 |
|---|---|---|---|---|---|---|
| S011 | 0.9459 | 0.0060 | 0.0476 | 0.0005 | 0.0000 | 0.0000 |
| S012 | 0.0415 | 0.8920 | 0.0017 | 0.0647 | 0.0000 | 0.0000 |
| S021 | 0.0614 | 0.0003 | 0.9361 | 0.0021 | 0.0000 | 0.0002 |
| S022 | 0.0021 | 0.0517 | 0.0095 | 0.9357 | 0.0009 | 0.0000 |
| S031 | 0.0007 | 0.0000 | 0.0000 | 0.0106 | 0.9774 | 0.0113 |
| S032 | 0.0045 | 0.0000 | 0.0403 | 0.0000 | 0.0336 | 0.9217 |
| Sum of diagonals: 5.608926 | | | | | | |
| Mean of diagonals: 0.934821 | | | | | | |
| Minimal of diagonals: 0.8920 | | | | | | |

with $S_0 = 500, S_c = 0, P_c = P_0 = 0.6$. The results are similar as shown by the high ARI values in Table 12. For $S_0$, since it is the minimal size of a cluster to be recognized as a meaningful one, results won't change as long as $S_0$ is less than the minimal size of the clusters that we get. For $P_0$, we get the same result for any $P_0$ within $(0.6, 0.75)$.

##### 3.6 Results from TRDG and MVCA

*Results from TRDG*

In this section, we show the result of applying TRDG to alanine dipeptide. We follow the steps in [5]: (1) In clustering step, microstates are obtained by grid clustering (see Section 3.2); (2) Estimating the free energy of each microstates $F_i \sim -kT\ln(Z_i)$ [5], with $Z_i$ being the number of times the system visited the microstate; (3) Estimating the energy barrier $F_{ij} \sim -kT\ln(Z_{ij})$ [5] between microstates, where $Z_{ij}$ is the minimum cut between microstates found by using Gomory-Hu algorithm implemented in the function "gomory_hu_tree" from the Python package "igraph"*; (4) Starting with the lowest energy barrier $F_{ij}$, we construct the disconnectivity graph by connecting successively microstates in order of increasing $F_{ij}$. This step clusters microstates by using hierarchical clustering with single linkage.

To better illustrate the behavior of TRDG on alanine dipeptide, we construct TRDG based on microstates from grids with different sizes, i.e, a 10×10 grid, a 20×20 grid and an 80×80 grid. Figure 7 shows the results. Figure 7(A) shows the benchmark clustering results [6] of microstates defined by a 10 × 10 grid. Note that ideally a 10×10 grid will give 100 clusters (microstates); however, some of them are empty, which result in 90 non-empty clusters (microstates) as shown in Figure 7(A). The numbers from 1 to 90 are the indexes of microstates, and their locations denote the centers of the corresponding microstates. Figure 7(B) shows the TRDG † results based on microstates defined in Figure 7(A). As shown in Figure 7(B), TRDG correctly recognized the microstates 31, 83, 75 and 72 as local stable states. However, microstates 72 and 83 should not be clustered separately since they are actually from the same cluster S1. In addition, we represent the TRDG results in Figure 7(B) as a phylogenetic tree shown in Figure 7(C), which contains the same clustering structure as the one in (B), and is easier to read with the label coloring. As shown in Figure 7(C), TRDG does not perform well on this dataset in terms of clustering, since the benchmark clusters are mixed up. Figure 7(D) and Figure 7(E) shows the TRDG results based on microstates defined by a 20 × 20 grid and an 80 × 80 grid, respectively. Since we care more about the performance of TRDG on clustering, we only show the phylogenetic tree representation of the TRDG for presenting clearly the cluster structure. By refining the microstates, the clustering structure from TRDG is not improved as shown in Figure 7(D & E).

*. https://igraph.org/python/

TABLE 9
Transition matrix between clusters of Alanine dipeptide obtained by PCCA+′

| | S1 | S2 | S3 | S4 | S5 |
|---|---|---|---|---|---|
| S1 | 0.9049 | 0.0616 | 0.0000 | 0.0000 | 0.0335 |
| S2 | 0.0257 | 0.9592 | 0.0113 | 0.0008 | 0.0030 |
| S3 | 0.0000 | 0.0005 | 0.9591 | 0.0257 | 0.0147 |
| S4 | 0.0000 | 0.0001 | 0.0668 | 0.5796 | 0.3535 |
| S5 | 0.0001 | 0.0000 | 0.0025 | 0.0241 | 0.9732 |
| Sum of diagonals: 4.376039 | | | | | |
| Mean of diagonals: 0.8752078 | | | | | |
| Minimal of diagonals: 0.5796 | | | | | |

TABLE 10
Transition matrix between clusters of Alanine dipeptide obtained by Gibbs′

| | S0 | S1 | S2 | S3 | S4 |
|---|---|---|---|---|---|
| S0 | 0.2867 | 0.1629 | 0.2686 | 0.2242 | 0.0576 |
| S1 | 0.0048 | 0.9417 | 0.0037 | 0.0013 | 0.0486 |
| S2 | 0.0256 | 0.0112 | 0.9497 | 0.0115 | 0.0019 |
| S3 | 0.3437 | 0.0608 | 0.1761 | 0.3414 | 0.0780 |
| S4 | 0.0021 | 0.0609 | 0.0008 | 0.0017 | 0.9345 |
| Sum of diagonals: 3.4540 | | | | | |
| Mean of diagonals: 0.6908 | | | | | |
| Minimal of diagonals: 0.2867 | | | | | |

Note that we compare different methods based on microstates defined by an 80 × 80 grid. According to the results in SI Figure 7(E) and Figure 4 in the main paper, TRDG did not performance well in terms of clustering.

*Results from MVCA*

In this section, we show the result of applying TRDG to alanine dipeptide. We follow the steps in [7]: (1) using symmetric Jensen-Shannon divergence to measure the similarity between microstates; (2) using agglomerative clustering with Wards minimum variance criterion to cluster microstates into macrostates (metastable states), and cutting the hierarchical clustering tree to give 6 clusters. The results are show in Figure 8.

†. The disconnectivity graph is plotted by using the package "disconnectionDPS" downloaded from http://www-wales.ch.cam.ac.uk/software.html

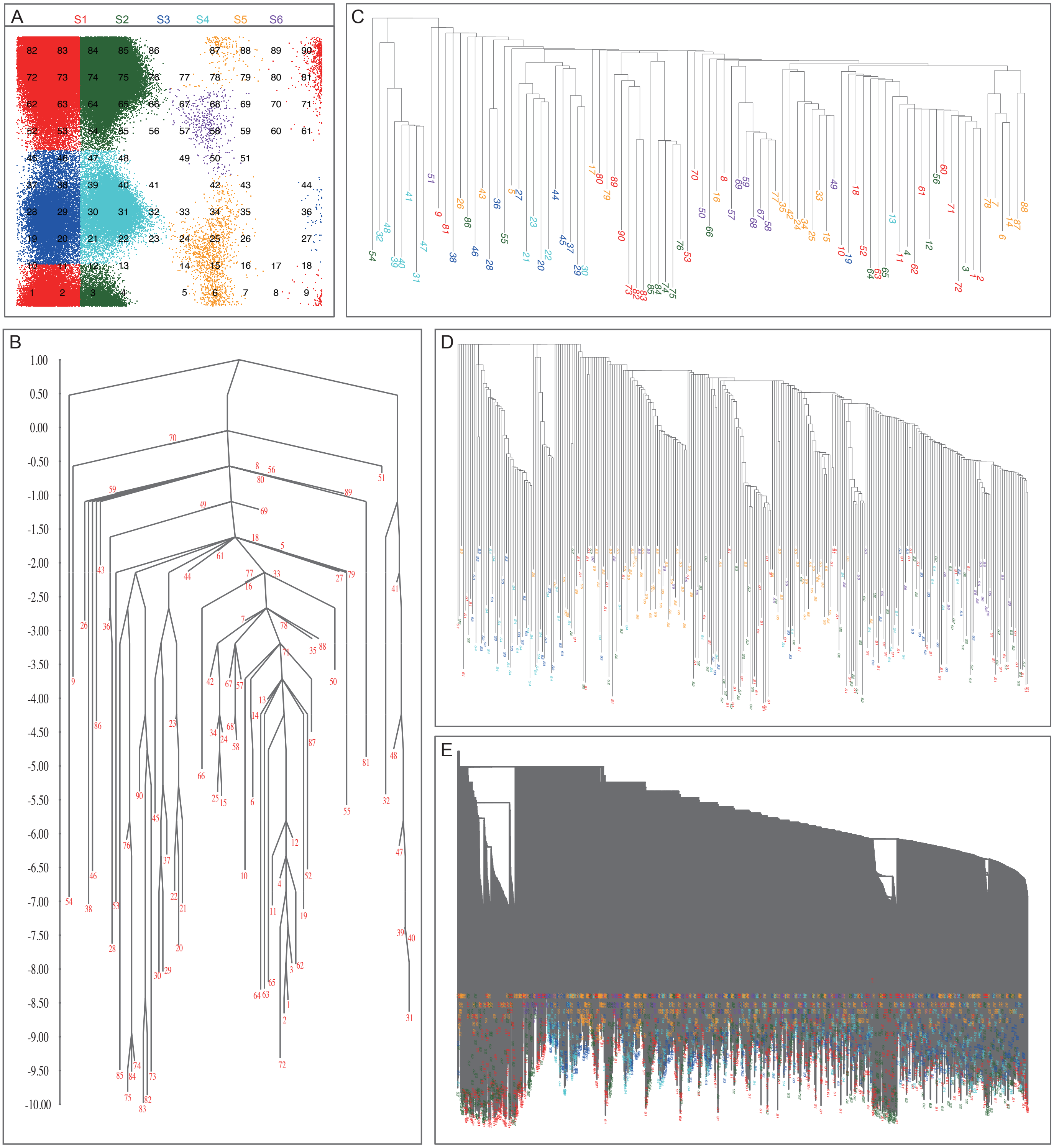

Fig. 7. Transition disconnectivity graph (TRDG) of alanine dipeptide. The y-axis for (B-E) shows the free energy from low to high. (A) A 10×10 grid generates 90 non-empty microstates from 6 benchmark clusters (S1-S6) with cluster color code at the top. The numbers located at the center of each microstate are the indexes of microstates. (B) TRDG results with microstates defined by (A). The label of each node corresponds to the index of the microstate in (A). (C) Phylogenetic tree representation of the TRDG results in (B). It contains the same information in (B), but with extra coloring for the labels. The label of each node corresponds to the index of the microstate in (A). The color of labels corresponds to the cluster color in (A). (D) Phylogenetic tree representation of the TRDG for microstates defined by a 20 × 20 grid. The color of labels corresponds to the cluster color in (A). (E) Phylogenetic tree representation of the TRDG for microstates defined by an 80 × 80 grid. The color of labels corresponds to the cluster color in (A).

### 4 Details about analyzing HP35 Nle/Nle

#### 4.1 The value of $d_0$

We applied CAPT on the MD data of HP35 Nle/Nle by using $P_0 = 0.7, P_c = 0.95, S_0 = 500, S_c = 10000$, and the Gaussian kernel. The partition tree is shown in Figure 9. We then tried 5 different values for $d_0$ to compute the LDc and

TABLE 11
Transition matrix between clusters of Alanine dipeptide obtained by Gibbs″

| | S0 | S1 | S2 | S3 | S4 | S5 | S6 |
|---|---|---|---|---|---|---|---|
| S0 | 0.9258 | 0.0055 | 0.0045 | 0.0030 | 0.0035 | 0.0562 | 0.0015 |
| S1 | 0.0015 | 0.9337 | 0.0021 | 0.0010 | 0.0607 | 0.0008 | 0.0003 |
| S2 | 0.1087 | 0.2027 | 0.2510 | 0.1638 | 0.0792 | 0.1329 | 0.0617 |
| S3 | 0.0502 | 0.0774 | 0.1363 | 0.2846 | 0.2388 | 0.0785 | 0.1341 |
| S4 | 0.0008 | 0.0477 | 0.0007 | 0.0024 | 0.9418 | 0.0062 | 0.0004 |
| S5 | 0.0670 | 0.0044 | 0.0072 | 0.0054 | 0.0416 | 0.8740 | 0.0004 |
| S6 | 0.0182 | 0.0166 | 0.0363 | 0.0979 | 0.0269 | 0.0047 | 0.7994 |
| Sum of diagonals: 5.0102 | | | | | | | |
| Mean of diagonals: 0.7157 | | | | | | | |
| Minimal of diagonals: 0.2510 | | | | | | | |

TABLE 12
ARI between cluster labels resulted from different kernel functions for analyzing Alanine dipeptide

| ARI | von Mises | Epanechnikov |
|---|---|---|
| Gaussian | 0.995622 | 0.9953416 |
| Epanechnikov | 0.9909818 | - |

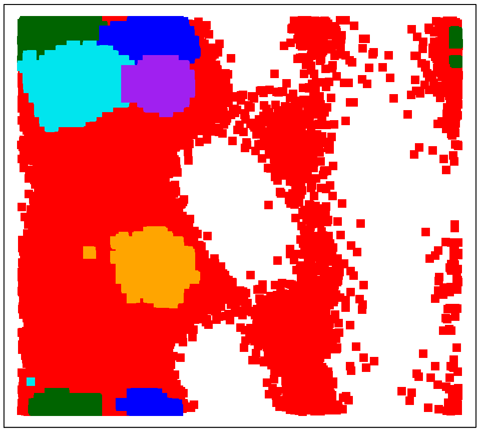

Fig. 8. Clustering results of alanine dipeptide from MVCA. Different colors present different clusters.

LDa values for the obtained clusters. Results are given in Table 13. We find that the order of the maximal LDc of these clusters do not vary with $d_0$. However, the stable structure, i.e., the frame with maximal LDc in each cluster, may vary with $d_0$, as shown in Figure 10. In the figure, the structures with the same color in each row are exactly the same frame. The MAD between the center of C39 under case B and case C is 0.1727754; The center of each cluster does not vary with $d_0$, when $d_0 \leq 0.1972243$, and we set $d_0 = 0.1757350$.

#### 4.2 Comparison with MPP

LDs of clusters obtained by MPP ‡ are given in Table 14. Corresponding structures with the maximal LDc in each cluster are given in Figure 11. We find that the LDc's of cluster U and N2 are 67 and 8 respectively when $d_0 = 0.1757350$, which are smaller than the 6-th most stable cluster found by CAPT (LDc=763).

‡. The clustering labels are provided by the authors of [3]

TABLE 13
The maximal local density LDc&LDa of each CAPT cluster calculated using different $d_0$. ST stands for similarity threshold, Q. is the short for quantile.

| ST / Q. | $d_0 =$ 0.3418513 (50%) | | $d_0 =$ 0.2318932 (5%) | | $d_0 =$ 0.1972243 (0.5%) | | $d_0 =$ 0.1757350 (0.05%) | | $d_0 =$ 0.1594225 (0.005%) | |
|---|---|---|---|---|---|---|---|---|---|---|
| | LDc | LDa | LDc | LDa | LDc | LDa | LDc | LDa | LDc | LDa |
| C1 | 12 | 12 | 3 | 3 | 2 | 2 | 1 | 1 | 1 | 1 |
| C2 | 46 | 46 | 3 | 3 | 1 | 1 | 1 | 1 | 1 | 1 |
| C3 | 67 | 67 | 4 | 4 | 1 | 1 | 1 | 1 | 1 | 1 |
| C4 | 22 | 23 | 2 | 2 | 1 | 1 | 1 | 1 | 1 | 1 |
| C5 | 24 | 24 | 3 | 3 | 1 | 1 | 1 | 1 | 1 | 1 |
| C6 | 36 | 36 | 4 | 4 | 1 | 1 | 1 | 1 | 1 | 1 |
| C7 | 33 | 33 | 3 | 3 | 2 | 2 | 1 | 1 | 1 | 1 |
| C8 | 48 | 48 | 4 | 4 | 1 | 1 | 1 | 1 | 1 | 1 |
| C9 | 180 | 220 | 42 | 42 | 6 | 6 | 3 | 3 | 1 | 1 |
| C10 | 22 | 22 | 2 | 2 | 2 | 2 | 1 | 1 | 1 | 1 |
| C11 | 15 | 15 | 3 | 3 | 1 | 1 | 1 | 1 | 1 | 1 |
| C12 | 28 | 28 | 3 | 3 | 2 | 2 | 1 | 1 | 1 | 1 |
| C13 | 23 | 23 | 7 | 7 | 2 | 2 | 1 | 1 | 1 | 1 |
| C14 | 19 | 19 | 3 | 3 | 1 | 1 | 1 | 1 | 1 | 1 |
| C15 | 46 | 46 | 3 | 3 | 1 | 1 | 1 | 1 | 1 | 1 |
| C16 | 35 | 35 | 5 | 5 | 2 | 2 | 2 | 2 | 1 | 1 |
| C17 | 552 | 1322 | 61 | 66 | 12 | 12 | 2 | 2 | 1 | 1 |
| C18 | 46 | 50 | 6 | 6 | 1 | 1 | 1 | 1 | 1 | 1 |
| C19 | 31 | 32 | 6 | 6 | 1 | 1 | 1 | 1 | 1 | 1 |
| C20 | 80 | 80 | 12 | 12 | 3 | 3 | 1 | 1 | 1 | 1 |
| C21 | 103 | 103 | 22 | 22 | 4 | 4 | 1 | 1 | 1 | 1 |
| C22 | 37 | 50 | 5 | 5 | 2 | 2 | 1 | 1 | 1 | 1 |
| C23 | 47 | 47 | 4 | 4 | 2 | 2 | 1 | 1 | 1 | 1 |
| C24 | 69 | 69 | 7 | 7 | 2 | 2 | 2 | 2 | 1 | 1 |
| C25 | 89 | 92 | 16 | 16 | 3 | 3 | 1 | 1 | 1 | 1 |
| C26 | 62 | 62 | 3 | 3 | 1 | 1 | 1 | 1 | 1 | 1 |
| C27 | 79 | 79 | 6 | 6 | 2 | 2 | 1 | 1 | 1 | 1 |
| C28 | 104 | 104 | 38 | 38 | 4 | 4 | 2 | 2 | 1 | 1 |
| C29 | 41 | 43 | 4 | 4 | 2 | 2 | 1 | 1 | 1 | 1 |
| C30 | 180 | 180 | 10 | 10 | 2 | 2 | 1 | 1 | 1 | 1 |
| C31 | 44 | 46 | 6 | 6 | 2 | 2 | 1 | 1 | 1 | 1 |
| C32 | 87 | 87 | 5 | 5 | 1 | 1 | 1 | 1 | 1 | 1 |
| C33 | 88 | 88 | 11 | 11 | 3 | 3 | 2 | 2 | 1 | 1 |
| C34 | 28 | 30 | 4 | 4 | 2 | 2 | 1 | 1 | 1 | 1 |
| C35 | 34 | 36 | 6 | 6 | 2 | 2 | 1 | 1 | 1 | 1 |
| C36 | 29 | 29 | 3 | 3 | 1 | 1 | 1 | 1 | 1 | 1 |
| C37 | 230 | 5977 | 25 | 25 | 5 | 5 | 3 | 3 | 1 | 1 |
| C38 | 63 | 63 | 4 | 4 | 2 | 2 | 1 | 1 | 1 | 1 |
| C39 | 2225 | 74916 | 627 | 638 | 126 | 126 | 21 | 21 | 4 | 4 |
| C40 | 126 | 144 | 10 | 10 | 3 | 3 | 1 | 1 | 1 | 1 |
| C41 | 113 | 121 | 10 | 10 | 3 | 3 | 1 | 1 | 1 | 1 |
| C42 | 66 | 66 | 7 | 7 | 2 | 2 | 1 | 1 | 1 | 1 |
| C43 | 109 | 109 | 9 | 9 | 3 | 3 | 2 | 2 | 1 | 1 |
| C44 | 59 | 59 | 9 | 9 | 2 | 2 | 2 | 2 | 1 | 1 |
| C45 | 45 | 46 | 6 | 6 | 2 | 2 | 1 | 1 | 1 | 1 |
| C46 | 140 | 140 | 40 | 40 | 4 | 4 | 2 | 2 | 1 | 1 |
| C47 | 184 | 135321 | 23 | 1228 | 5 | 20 | 2 | 2 | 1 | 1 |
| C48 | 197 | 200 | 23 | 23 | 6 | 6 | 2 | 2 | 1 | 1 |
| C49 | 88 | 89 | 8 | 8 | 2 | 2 | 1 | 1 | 1 | 1 |
| C50 | 325 | 140438 | 20 | 20 | 5 | 5 | 2 | 2 | 1 | 1 |
| C51 | 167 | 169 | 12 | 12 | 3 | 3 | 2 | 2 | 1 | 1 |
| C52 | 158 | 180 | 21 | 21 | 6 | 6 | 2 | 2 | 1 | 1 |
| C53 | 4283 | 7914 | 554 | 573 | 83 | 83 | 14 | 14 | 3 | 3 |
| C54 | 90 | 118 | 8 | 9 | 2 | 2 | 1 | 1 | 1 | 1 |
| C55 | 104 | 208 | 34 | 34 | 11 | 11 | 3 | 3 | 2 | 2 |
| C56 | 5504 | 6852 | 554 | 554 | 110 | 110 | 20 | 20 | 4 | 4 |
| C57 | 1719 | 2734 | 62 | 63 | 7 | 7 | 2 | 2 | 1 | 1 |
| C58 | 4735 | 14087 | 118 | 118 | 38 | 38 | 14 | 14 | 4 | 4 |
| C59 | 306331 | 307503 | 86021 | 86022 | 28297 | 28299 | 6854 | 6854 | 1303 | 1303 |

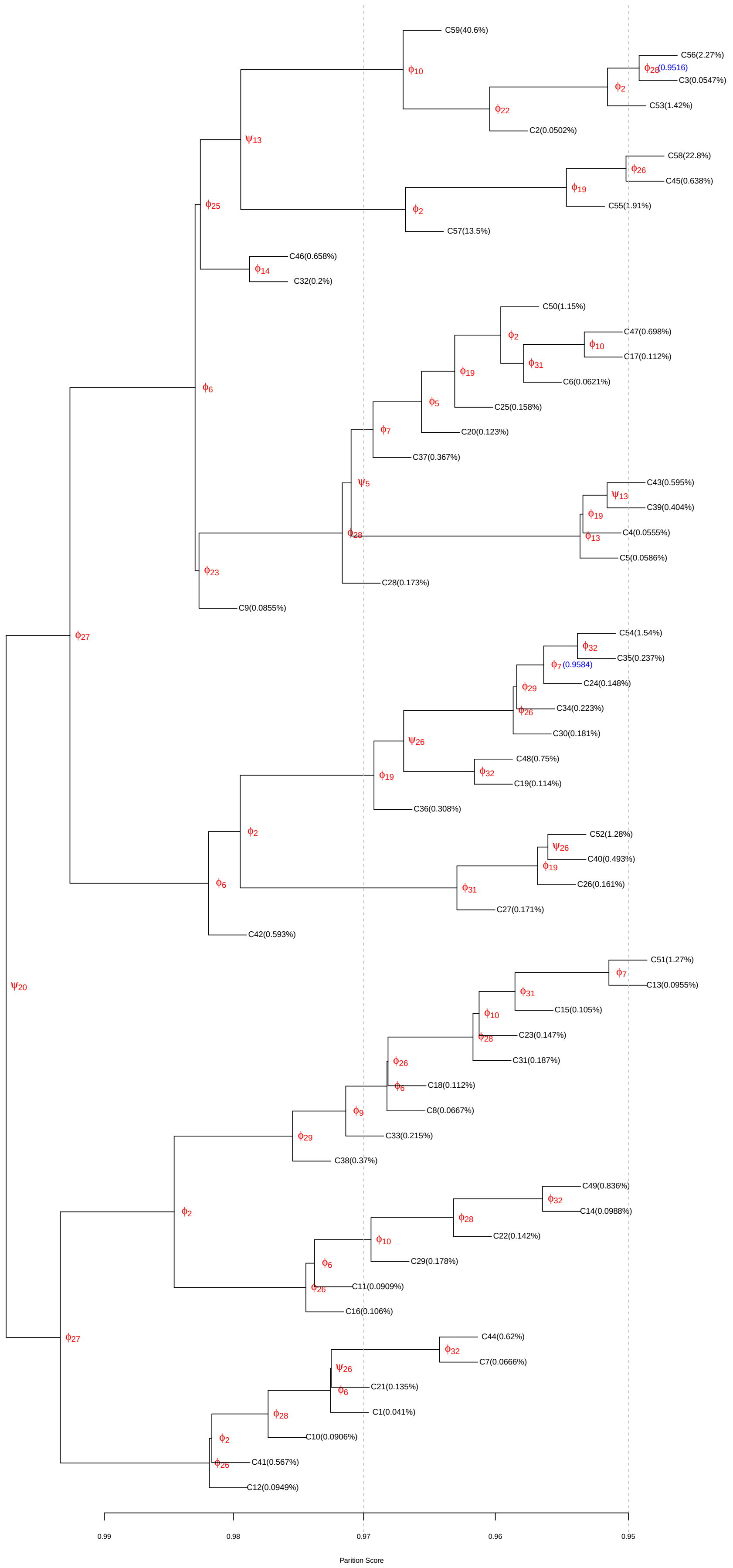


Fig. 9. Partition tree of HP35 Nle/Nle, with $S_0 = 500, P_0 = 0.7, S_c = 10000, P_c = 0.95$ and the Gaussian kernel.

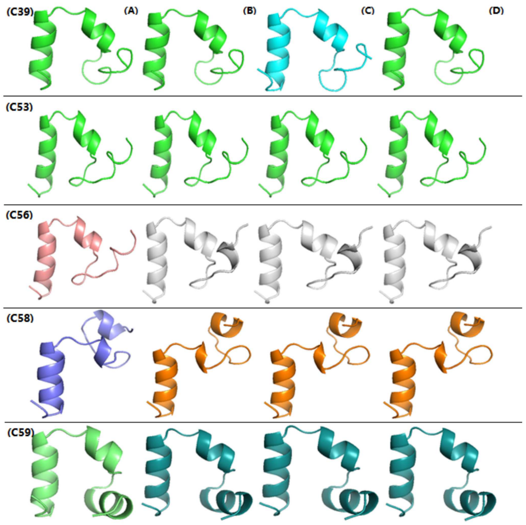


Fig. 10. Different stable structures resulting from different $d_0$ by CAPT. Each column corresponds to different $d_0$: (A) $d_0 = 0.2318932$; (B) $d_0 = 0.1972243$; (C) $d_0 = 0.1757350$; (D) $d_0 = 0.1594225$. The MAD between the centers of C39 under Case B and Case C is 0.1727754. The structures with the same color in each row are exactly the same frame.

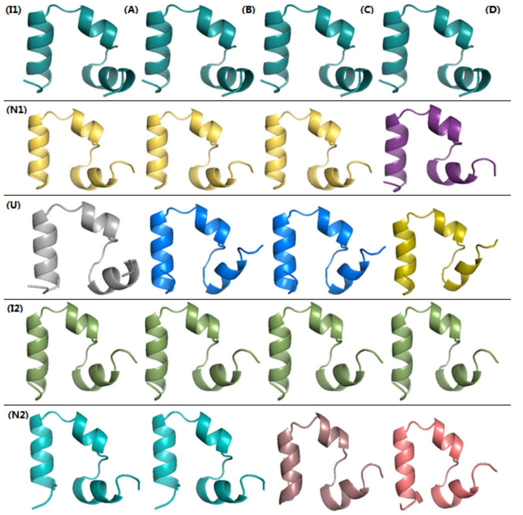


Fig. 11. Different stable structures resulting from different $d_0$ by MPP. MPP found five clusters: I1, I2, U, N1 and N2. Each column corresponds to different $d_0$: (A) $d_0 = 0.2318932$; (B) $d_0 = 0.1972243$; (C) $d_0 = 0.1757350$; (D) $d_0 = 0.1594225$. Detailed values of LDc and LDa are given in Table 14. The structures with the same color in each row are exactly the same frame.

##### 4.3 Sensitivity analysis

In this part, we conduct the sensitivity analysis for the parameters $S_0$ and $P_0$, as well as the kernel function for density estimation. $S_0$, as discussed before, only determines the minimal size of each cluster we obtained. We set $S_0 = 10000$ for previous experiments. Here we also give the results while using $S_0 = 500$ with $P_0 = 0.7$ and the Gaussian kernel. Since there are many clusters, we first set $P_c = 0.95$. The results are given in Table 13 and the partition tree is shown in Figure 9. These results show that there is only one stable cluster C59 with LDc=6854. We then set $P_c = 0.7$ and run CAPT on C59. The partition tree is shown in Figure 13, where we find 6 stable clusters, as shown in Table 15. These are exactly the same results found by setting $S_0 = 10000$. However, with a too large $S_0$, we can not get the right number of clusters. Taking Alanine dipeptide as an example, we only get 4 clusters if we set $S_0 = 1948$ (i.e., 1% of the whole population), because there are only 1529 and 425 frames in the clusters S5 and S6 of Figure 4(A) in the main paper, respectively. So if one wants to explore the whole cluster structure, one should set a smaller value for $S_0$, say, $100 \leq S_0 \leq 500$. For complex biomolecules, one may be interested only in the stable structures, and a larger value for $S_0$ seems more efficient in such cases.

TABLE 14
The maximal local density LDc&LDa of each MPP cluster calculated using different $d_0$. ST stands for similarity threshold, Q. is the short for quantile.

| ST/Q. | | I1 | N1 | U | I2 | N2 |
|---|---|---|---|---|---|---|
| $d_0 = 0.3418513$ | LDc | 114766 | 111358 | 10134 | 84735 | 10647 |
| (50%) | LDa | 307002 | 294150 | 119336 | 297628 | 44064 |
| $d_0 = 0.2318932$ | LDc | 71943 | 55576 | 1978 | 20080 | 499 |
| (5%) | LDa | 82447 | 58818 | 2223 | 31423 | 535 |
| $d_0 = 0.1972243$ | LDc | 27375 | 15659 | 376 | 4356 | 49 |
| (0.5%) | LDa | 28299 | 15726 | 379 | 4759 | 49 |
| $d_0 = 0.1757350$ | LDc | 6766 | 3789 | 67 | 816 | 8 |
| (0.05%) | LDa | 6854 | 3790 | 67 | 839 | 8 |
| $d_0 = 0.1594225$ | LDc | 1293 | 737 | 12 | 130 | 3 |
| (0.005%) | LDa | 1303 | 737 | 12 | 133 | 3 |

TABLE 15
LDc of HP35 Nle/Nle clusters in the partition tree shown in Figure 13. $d_0$ is set as 0.1757350.

| Cluster | L1 | L2 | L3 | L4 | L5 | L6 | L7 | L8 | L9 |
|---|---|---|---|---|---|---|---|---|---|
| LDc | 1 | 1 | 1 | 2 | 1 | 1 | 1 | 8 | 1 |
| Cluster | L10 | L11 | L12 | L13 | L14 | L15 | L16 | L17 | L18 |
| LDc | 1 | 3 | 2 | 1 | 1 | 14 | 2 | 1 | 2 |
| Cluster | L19 | L20 | L21 | L22 | L23 | L24 | L25 | L26 | L27 |
| LDc | 1 | 1 | 1 | 1 | 1 | 5 | 1 | 1 | 1 |
| Cluster | L28 | L29 | L30 | L31 | L32 | L33 | L34 | L35 | L36 |
| LDc | 2 | 1 | 1 | 1 | 22 | 7 | 1 | 1 | 3 |
| Cluster | L37 | L38 | L39 | L40 | L41 | L42 | L43 | L44 | L45 |
| LDc | 1 | 3 | 2 | 2 | 1 | 2 | 1 | 3 | 1 |
| Cluster | L46 | L47 | L48 | L49 | L50 | L51 | L52 | L53 | L54 |
| LDc | 2 | 1 | 2 | 1 | 1 | 2 | 1 | 2 | 7 |
| Cluster | L55 | L56 | L57 | L58 | L59 | L60 | L61 | L62 | L63 |
| LDc | 2 | 4 | 1 | 2 | 11 | 7 | 11 | 1 | 2 |
| Cluster | L64 | L65 | L66 | L67 | L68 | L69 | L70 | L71 | L72 |
| LDc | 2 | 4 | 2 | 5 | 8 | 3 | 2 | 5 | 4 |
| Cluster | L73 | L74 | L75 | L76 | L77 | L78 | L79 | L80 | L81 |
| LDc | 2 | 2 | 10 | 20 | 5 | 2 | 2 | 3 | 3 |
| Cluster | L82 | L83 | L84 | L85 | L86 | L87 | L88 | L89 | L90 |
| LDc | 8 | 2 | 67 | 3 | 51 | 12 | 3167 | 1321 | 763 |
| Cluster | L91 | L92 | L93 | | | | | | |
| LDc | 834 | 3662 | 6854 | | | | | | |

For $P_0$, we tried five different values {0.6, 0.65, 0.7, 0.75, 0.8}, with $S_0 = 10000$ and the Gaussian kernel. The resulting partition trees are given in Figure 14, Figure 15, Figure 16, Figure 17 and Figure 18. The corresponding LDc values are given in Table 17, Table 18, Table 19, Table 20 and Table 21. A summary of stable clusters is given in Table 22. There are only 5 stable clusters when $P_0 = 0.8$. For $0.6 \leq P_0 \leq 0.75$, we get 6 stable clusters. Interestingly, although

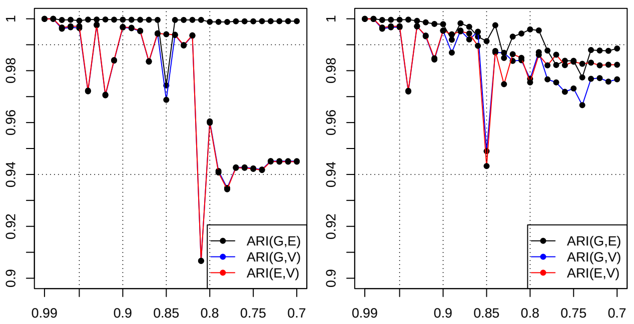


Fig. 12. ARI between cluster labels based on different kernel functions under different value of $S_0$ with $S_c = 10000$, $P_c = P_0 = 0.7$. The left panel is for the case $S_0 = 10000$, and the right panel is for the case $S_0 = 500$. In the figure, the x-axis is the energy barrier and the y-axis is the ARI. ARI(E,V) is the ARI between cluster labels based on the Epanechnikov kernel and the von Mises kernel, ARI(G,E) is the ARI between cluster labels based on the Gaussian kernel and the Epanechnikov kernel, ARI(G,V) is the ARI between cluster labels based on the Gaussian kernel and the von Mises kernel.

the resulting partition trees are different for different values of $P_0$, the key energy barriers between these 6 stable clusters are the same.

TABLE 16
Transition Matrix between meaningful clusters obtained by CAPT for HP35 Nle/Nle. 'Other' are conformations not belonging to S1-S6.

| | other | S1 | S2 | S3 | S4 | S5 | S6 |
|---|---|---|---|---|---|---|---|
| other | 0.9886 | 0.0003 | 0.0033 | 0.0007 | 0.0027 | 0.0002 | 0.0041 |
| S1 | 0.0101 | 0.9230 | 0.0006 | 0.0062 | 0.0000 | 0.0600 | 0.0001 |
| S2 | 0.0662 | 0.0004 | 0.9199 | 0.0000 | 0.0090 | 0.0000 | 0.0046 |
| S3 | 0.0111 | 0.0040 | 0.0000 | 0.9099 | 0.0005 | 0.0736 | 0.0009 |
| S4 | 0.0551 | 0.0000 | 0.0093 | 0.0008 | 0.8826 | 0.0001 | 0.0520 |
| S5 | 0.0067 | 0.0641 | 0.0000 | 0.1430 | 0.0003 | 0.7827 | 0.0032 |
| S6 | 0.0361 | 0.0000 | 0.0020 | 0.0004 | 0.0223 | 0.0009 | 0.9383 |

For the kernel function used for density estimation, we tried three different kernels: Gaussian, von Mises and Epanechnikov. They result in the same stable clusters, as shown in Table 23 and Table 24. Note that for different kernel functions, the partition trees are a bit different, as shown in Figure 19 and Figure 20. In addition, we show the ARI values between cluster labels from different kernel functions in Figure 12. As we can see from the figure, ARI is sensitive to the partition score cutoff, which is reasonable. Different kernel functions result in density estimates of different smoothness. ARI may drop at some cutoff, however it will be pulled up later as shown in the figure. For example, when cutoff=0.82, ARI(G,V)> 0.99; however, it drops to 0.9 when cutoff=0.81, and goes to 0.96 when cutoff=0.8. Even for the same angle, the partition scores are different under different kernel functions. That is, from a local point of view, the result is sensitive to the kernel function; however, from a global point of view, it is insensitive to the kernel function.

TABLE 17
LDc with $d_0 = 0.1757350$ of centers of HP35 Nle/Nle in partition tree shown in Figure 14

| | | | | | | | | | |
|---|---|---|---|---|---|---|---|---|---|
| Cluster | L1 | L2 | L3 | L4 | L5 | L6 | L7 | L8 | L9 |
| LDc | 2 | 1 | 2 | 3 | 2 | 2 | 2 | 1 | 12 |
| Cluster | L10 | L11 | L12 | L13 | L14 | L15 | L16 | L17 | L18 |
| LDc | 1 | 14 | 4 | 2 | 3 | 2 | 1 | 2 | 2 |
| Cluster | L19 | L20 | L21 | L22 | L23 | L24 | L25 | L26 | L27 |
| LDc | 1 | 2 | 3 | 51 | 2 | 2 | 1 | 2 | 1 |
| Cluster | L28 | L29 | L30 | L31 | L32 | L33 | L34 | L35 | L36 |
| LDc | 2 | 8 | 2 | 2 | 1 | 2 | 1 | 1 | 1 |
| Cluster | L37 | L38 | L39 | L40 | L41 | L42 | L43 | L44 | L45 |
| LDc | 4 | 1 | 1 | 2 | 2 | 1 | 2 | 2 | 2 |
| Cluster | L46 | L47 | L48 | L49 | L50 | L51 | L52 | L53 | L54 |
| LDc | 1 | 2 | 2 | 22 | 2 | 3 | 2 | 2 | 1 |
| Cluster | L55 | L56 | L57 | L58 | L59 | L60 | L61 | L62 | L63 |
| LDc | 2 | 3 | 5 | 3 | 3 | 8 | 2 | 2 | 2 |
| Cluster | L64 | L65 | L66 | L67 | L68 | L69 | L70 | L71 | L72 |
| LDc | 1 | 2 | 2 | 4 | 3 | 2 | 2 | 2 | 3 |
| Cluster | L73 | L74 | L75 | L76 | L77 | L78 | L79 | L80 | L81 |
| LDc | 21 | 2 | 2 | 14 | 20 | 2 | 2 | 2 | 2 |
| Cluster | L82 | L83 | L84 | L85 | L86 | L87 | L88 | L89 | |
| LDc | 3 | 1 | 1321 | 3167 | 834 | 763 | 3663 | 6854 | |

TABLE 18
LDc with $d_0 = 0.1757350$ of centers of HP35 Nle/Nle in the partition tree shown in Figure 15

| | | | | | | | | | |
|---|---|---|---|---|---|---|---|---|---|
| Cluster | L1 | L2 | L3 | L4 | L5 | L6 | L7 | L8 | L9 |
| LDc | 3 | 2 | 2 | 2 | 1 | 14 | 4 | 1 | 14 |
| Cluster | L10 | L11 | L12 | L13 | L14 | L15 | L16 | L17 | L18 |
| LDc | 2 | 3 | 2 | 1 | 2 | 2 | 2 | 2 | 1 |
| Cluster | L19 | L20 | L21 | L22 | L23 | L24 | L25 | L26 | L27 |
| LDc | 2 | 3 | 51 | 2 | 1 | 1 | 2 | 2 | 2 |
| Cluster | L28 | L29 | L30 | L31 | L32 | L33 | L34 | L35 | L36 |
| LDc | 2 | 2 | 2 | 8 | 3 | 2 | 2 | 2 | 4 |
| Cluster | L37 | L38 | L39 | L40 | L41 | L42 | L43 | L44 | L45 |
| LDc | 1 | 1 | 2 | 2 | 2 | 1 | 2 | 2 | 2 |
| Cluster | L46 | L47 | L48 | L49 | L50 | L51 | L52 | L53 | L54 |
| LDc | 1 | 2 | 2 | 1 | 1 | 2 | 2 | 3 | 1 |
| Cluster | L55 | L56 | L57 | L58 | L59 | L60 | L61 | L62 | L63 |
| LDc | 3 | 2 | 3 | 1 | 2 | 1 | 2 | 2 | 2 |
| Cluster | L64 | L65 | L66 | L67 | L68 | L69 | L70 | L71 | L72 |
| LDc | 2 | 2 | 4 | 3 | 3 | 1 | 5 | 3 | 2 |
| Cluster | L73 | L74 | L75 | L76 | L77 | L78 | L79 | L80 | L81 |
| LDc | 21 | 2 | 2 | 2 | 2 | 2 | 20 | 1 | 2 |
| Cluster | L82 | L83 | L84 | L85 | L86 | L87 | L88 | | |
| LDc | 12 | 1321 | 3167 | 834 | 763 | 3663 | 6854 | | |

#### 5 Remark on the von Mises kernel

Assuming that $\{\theta_i, i = 1, ...n\}$ follows vM$(\mu, \kappa)$, the density estimate using a standard von Mises kernel [8] is given by

$$\hat{f}(\theta, v) = \frac{1}{n(2\pi)I_0(v)} \sum_{i=1}^{n} e^{v\cos(\theta-\theta_i)}, \tag{1}$$

where

$$v = \{\frac{3n\hat{\kappa}^2 I_2(2\hat{\kappa})}{4\sqrt{\pi}I_0(\hat{\kappa})^2}\}^{\frac{2}{5}}, \tag{2}$$

$\hat{\kappa}$ is the MLE of $\kappa$, $I_r(v)$ is the modified Bessel function of order $r$, and $v$ takes the role of (inverse of) smoothing parameter. In practice, the computer gives $I_0(v) = \infty$ when $v > 709$, i.e., larger than the maximal number that the computer can handle with. Thus we set $v = 709$ when $v$ is actually bigger than 709 in analysis.

TABLE 19
LDc with $d_0 = 0.1757350$ of centers of HP35 Nle/Nle in the partition tree shown in Figure 16

| Cluster | L1 | L2 | L3 | L4 | L5 | L6 | L7 | L8 | L9 |
|---|---|---|---|---|---|---|---|---|---|
| LDc | 1 | 2 | 1 | 2 | 2 | 2 | 14 | 4 | 1 |
| Cluster | L10 | L11 | L12 | L13 | L14 | L15 | L16 | L17 | L18 |
| LDc | 3 | 2 | 2 | 2 | 14 | 2 | 3 | 51 | 1 |
| Cluster | L19 | L20 | L21 | L22 | L23 | L24 | L25 | L26 | L27 |
| LDc | 1 | 1 | 2 | 8 | 2 | 2 | 2 | 2 | 2 |
| Cluster | L28 | L29 | L30 | L31 | L32 | L33 | L34 | L35 | L36 |
| LDc | 2 | 2 | 2 | 4 | 1 | 1 | 2 | 2 | 2 |
| Cluster | L37 | L38 | L39 | L40 | L41 | L42 | L43 | L44 | L45 |
| LDc | 1 | 3 | 2 | 1 | 2 | 2 | 1 | 2 | 3 |
| Cluster | L46 | L47 | L48 | L49 | L50 | L51 | L52 | L53 | L54 |
| LDc | 2 | 1 | 2 | 2 | 3 | 1 | 2 | 2 | 1 |
| Cluster | L55 | L56 | L57 | L58 | L59 | L60 | L61 | L62 | L63 |
| LDc | 2 | 2 | 2 | 2 | 4 | 3 | 2 | 1 | 5 |
| Cluster | L64 | L65 | L66 | L67 | L68 | L69 | L70 | L71 | L72 |
| LDc | 1 | 2 | 21 | 3 | 1 | 2 | 2 | 2 | 2 |
| Cluster | L73 | L74 | L75 | L76 | L77 | L78 | L79 | L80 | L81 |
| LDc | 2 | 20 | 2 | 2 | 2 | 3 | 12 | 1321 | 3167 |
| Cluster | L82 | L83 | L84 | L85 | | | | | |
| LDc | 834 | 763 | 3663 | 6854 | | | | | |

TABLE 20
LDc with $d_0 = 0.1757350$ of centers of HP35 Nle/Nle in the partition tree shown in Figure 17

| Cluster | L1 | L2 | L3 | L4 | L5 | L6 | L7 | L8 | L9 |
|---|---|---|---|---|---|---|---|---|---|
| LDc | 1 | 2 | 1 | 2 | 51 | 2 | 14 | 1 | 3 |
| Cluster | L10 | L11 | L12 | L13 | L14 | L15 | L16 | L17 | L18 |
| LDc | 2 | 2 | 14 | 2 | 2 | 1 | 1 | 1 | 2 |
| Cluster | L19 | L20 | L21 | L22 | L23 | L24 | L25 | L26 | L27 |
| LDc | 8 | 2 | 2 | 2 | 2 | 2 | 2 | 2 | 1 |
| Cluster | L28 | L29 | L30 | L31 | L32 | L33 | L34 | L35 | L36 |
| LDc | 1 | 2 | 2 | 2 | 1 | 3 | 2 | 1 | 2 |
| Cluster | L37 | L38 | L39 | L40 | L41 | L42 | L43 | L44 | L45 |
| LDc | 2 | 1 | 2 | 3 | 2 | 1 | 2 | 3 | 1 |
| Cluster | L46 | L47 | L48 | L49 | L50 | L51 | L52 | L53 | L54 |
| LDc | 2 | 2 | 2 | 3 | 1 | 2 | 2 | 2 | 4 |
| Cluster | L55 | L56 | L57 | L58 | L59 | L60 | L61 | L62 | L63 |
| LDc | 2 | 2 | 2 | 1 | 1 | 2 | 21 | 3 | 1 |
| Cluster | L64 | L65 | L66 | L67 | L68 | L69 | L70 | L71 | L72 |
| LDc | 2 | 4 | 5 | 2 | 2 | 2 | 20 | 2 | 2 |
| Cluster | L73 | L74 | L75 | L76 | L77 | L78 | L79 | L80 | L81 |
| LDc | 2 | 2 | 5 | 3 | 12 | 1321 | 3167 | 834 | 763 |
| Cluster | L82 | L83 | | | | | | | |
| LDc | 3666 | 6854 | | | | | | | |

TABLE 21
LDc with $d_0 = 0.1757350$ of centers of HP35 Nle/Nle in the partition tree shown in Figure 18

| Cluster | L1 | L2 | L3 | L4 | L5 | L6 | L7 | L8 | L9 |
|---|---|---|---|---|---|---|---|---|---|
| LDc | 1 | 2 | 1 | 14 | 2 | 2 | 2 | 2 | 21 |
| Cluster | L10 | L11 | L12 | L13 | L14 | L15 | L16 | L17 | L18 |
| LDc | 2 | 2 | 1 | 1 | 2 | 2 | 2 | 2 | 3 |
| Cluster | L19 | L20 | L21 | L22 | L23 | L24 | L25 | L26 | L27 |
| LDc | 1 | 1 | 1 | 2 | 2 | 1 | 4 | 2 | 3 |
| Cluster | L28 | L29 | L30 | L31 | L32 | L33 | L34 | L35 | L36 |
| LDc | 3 | 2 | 2 | 1 | 3 | 1 | 2 | 1 | 2 |
| Cluster | L37 | L38 | L39 | L40 | L41 | L42 | L43 | L44 | L45 |
| LDc | 2 | 2 | 3 | 2 | 1 | 2 | 3 | 2 | 2 |
| Cluster | L46 | L47 | L48 | L49 | L50 | L51 | L52 | L53 | L54 |
| LDc | 3 | 2 | 2 | 4 | 20 | 2 | 2 | 2 | 1 |
| Cluster | L55 | L56 | L57 | L58 | L59 | L60 | L61 | L62 | L63 |
| LDc | 14 | 1 | 3 | 2 | 2 | 2 | 3 | 2 | 5 |
| Cluster | L64 | L65 | L66 | L67 | L68 | L69 | L70 | L71 | L72 |
| LDc | 2 | 2 | 12 | 3167 | 2 | 763 | 834 | 3764 | 6854 |

TABLE 22
clusters with maximal LDc (Top 6) under different $P_0$, where S1-6 is the cluster name used to give a unified comparison. Corresponding structure is given in Figure 6.

| | | S1 | S2 | S3 | S4 | S5 | S6 |
|---|---|---|---|---|---|---|---|
| $P_0 = 0.8$ | cluster | L67 | L69 | L71 | L70 | - | L72 |
| | LDc | 3167 | 763 | 3764 | 834 | - | 6854 |
| $P_0 = 0.75$ | cluster | L79 | L81 | L82 | L80 | L78 | L83 |
| | LDc | 3167 | 763 | 3666 | 834 | 1321 | 6854 |
| $P_0 = 0.7$ | cluster | L81 | L83 | L84 | L82 | L80 | L85 |
| | LDc | 3167 | 763 | 3663 | 834 | 1321 | 6854 |
| $P_0 = 0.65$ | cluster | L84 | L86 | L87 | L85 | L83 | L88 |
| | LDc | 3167 | 763 | 3663 | 834 | 1321 | 6854 |
| $P_0 = 0.6$ | cluster | L85 | L87 | L88 | L86 | L84 | L89 |
| | LDc | 3167 | 763 | 3663 | 834 | 1321 | 6854 |

TABLE 23
LDc with $d_0 = 0.1757350$ of centers of HP35 Nle/Nle in the partition tree shown in Figure 19

| Cluster | L1 | L2 | L3 | L4 | L5 | L6 | L7 | L8 | L9 |
|---|---|---|---|---|---|---|---|---|---|
| LDc | 1 | 2 | 1 | 2 | 2 | 2 | 14 | 4 | 1 |
| Cluster | L10 | L11 | L12 | L13 | L14 | L15 | L16 | L17 | L18 |
| LDc | 2 | 3 | 2 | 2 | 2 | 14 | 51 | 3 | 1 |
| Cluster | L19 | L20 | L21 | L22 | L23 | L24 | L25 | L26 | L27 |
| LDc | 1 | 1 | 2 | 8 | 2 | 2 | 2 | 2 | 2 |
| Cluster | L28 | L29 | L30 | L31 | L32 | L33 | L34 | L35 | L36 |
| LDc | 2 | 2 | 2 | 4 | 1 | 1 | 2 | 2 | 2 |
| Cluster | L37 | L38 | L39 | L40 | L41 | L42 | L43 | L44 | L45 |
| LDc | 1 | 3 | 2 | 1 | 2 | 2 | 1 | 2 | 3 |
| Cluster | L46 | L47 | L48 | L49 | L50 | L51 | L52 | L53 | L54 |
| LDc | 2 | 1 | 2 | 2 | 3 | 1 | 2 | 2 | 1 |
| Cluster | L55 | L56 | L57 | L58 | L59 | L60 | L61 | L62 | L63 |
| LDc | 2 | 2 | 2 | 2 | 4 | 3 | 2 | 1 | 5 |
| Cluster | L64 | L65 | L66 | L67 | L68 | L69 | L70 | L71 | L72 |
| LDc | 1 | 2 | 21 | 3 | 1 | 2 | 2 | 2 | 2 |
| Cluster | L73 | L74 | L75 | L76 | L77 | L78 | L79 | L80 | L81 |
| LDc | 2 | 20 | 2 | 2 | 2 | 3 | 12 | 1321 | 3167 |
| Cluster | L82 | L83 | L84 | L85 | | | | | |
| LDc | 834 | 763 | 3663 | 6854 | | | | | |

#### 6 Partition trees of HP35 Nle/Nle under different settings

TABLE 24
LDc with $d_0 = 0.1757350$ of centers of HP35 Nle/Nle in the partition tree shown in Figure 20

| | | | | | | | | | |
|---|---|---|---|---|---|---|---|---|---|
| Cluster | L1 | L2 | L3 | L4 | L5 | L6 | L7 | L8 | L9 |
| LDc | 1 | 1 | 2 | 2 | 1 | 2 | 4 | 1 | 14 |
| Cluster | L10 | L11 | L12 | L13 | L14 | L15 | L16 | L17 | L18 |
| LDc | 2 | 2 | 3 | 2 | 1 | 14 | 2 | 5 | 1 |
| Cluster | L19 | L20 | L21 | L22 | L23 | L24 | L25 | L26 | L27 |
| LDc | 3 | 2 | 2 | 1 | 2 | 2 | 1 | 8 | 2 |
| Cluster | L28 | L29 | L30 | L31 | L32 | L33 | L34 | L35 | L36 |
| LDc | 2 | 1 | 2 | 2 | 2 | 4 | 1 | 2 | 2 |
| Cluster | L37 | L38 | L39 | L40 | L41 | L42 | L43 | L44 | L45 |
| LDc | 2 | 3 | 1 | 2 | 2 | 2 | 1 | 2 | 2 |
| Cluster | L46 | L47 | L48 | L49 | L50 | L51 | L52 | L53 | L54 |
| LDc | 3 | 2 | 1 | 2 | 1 | 2 | 3 | 2 | 2 |
| Cluster | L55 | L56 | L57 | L58 | L59 | L60 | L61 | L62 | L63 |
| LDc | 51 | 1 | 2 | 2 | 2 | 4 | 2 | 2 | 3 |
| Cluster | L64 | L65 | L66 | L67 | L68 | L69 | L70 | L71 | L72 |
| LDc | 1 | 5 | 1 | 21 | 3 | 2 | 1 | 2 | 2 |
| Cluster | L73 | L74 | L75 | L76 | L77 | L78 | L79 | L80 | L81 |
| LDc | 2 | 2 | 2 | 2 | 2 | 12 | 20 | 2 | 3 |
| Cluster | L82 | L83 | L84 | L85 | L86 | L87 | | | |
| LDc | 1321 | 3167 | 834 | 3662 | 763 | 6854 | | | |

[8] Charles C Taylor. Automatic bandwidth selection for circular density estimation. *Computational Statistics & Data Analysis*, 52(7):3493–3500, 2008.

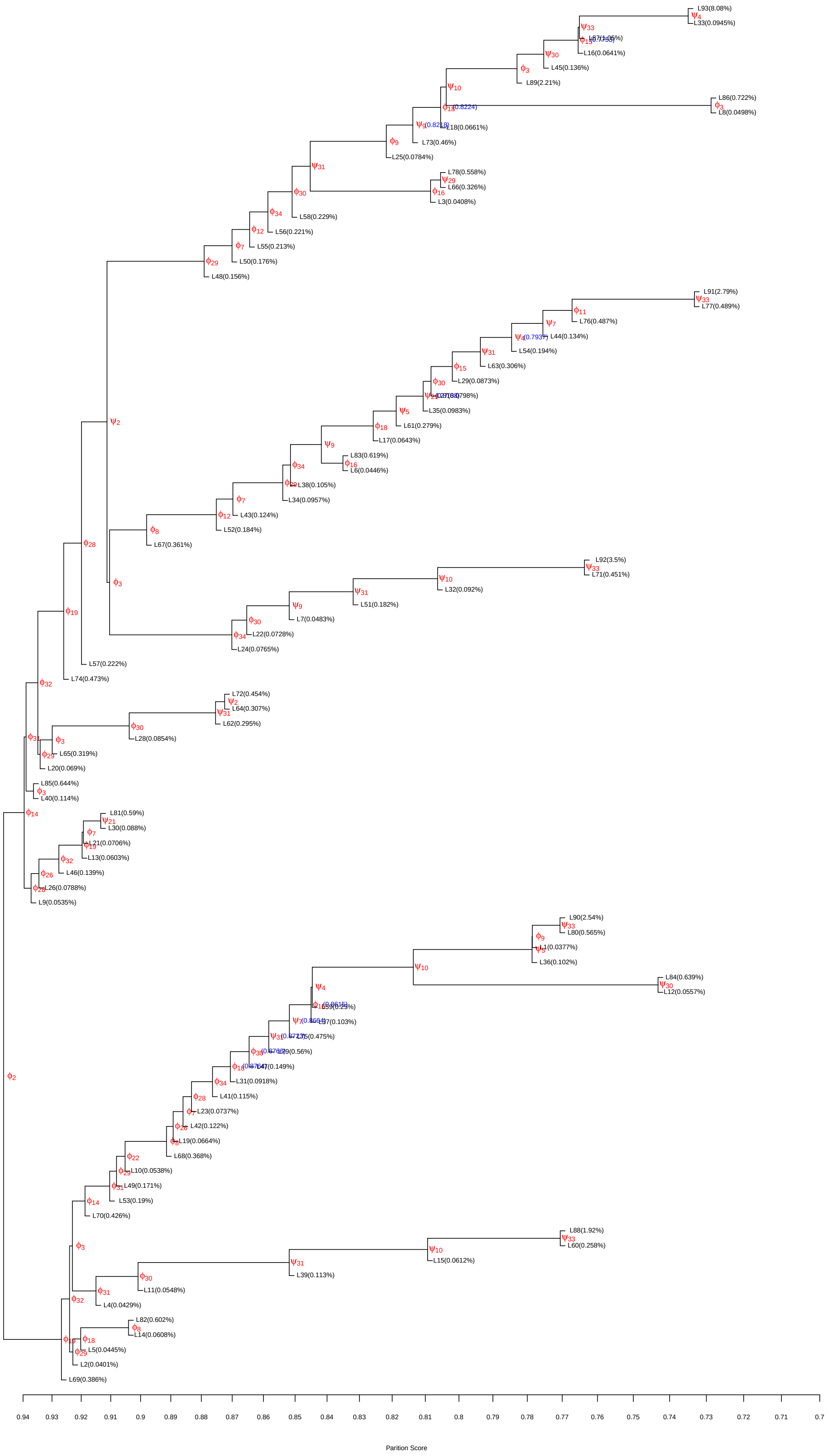


Fig. 13. Sub-partition tree of C59 in Figure 9 for HP35 Nle/Nle with $S_0 = 500$, $P_c = P_0 = 0.7$, and the Gaussian kernel. In the tree, the root node is C59.

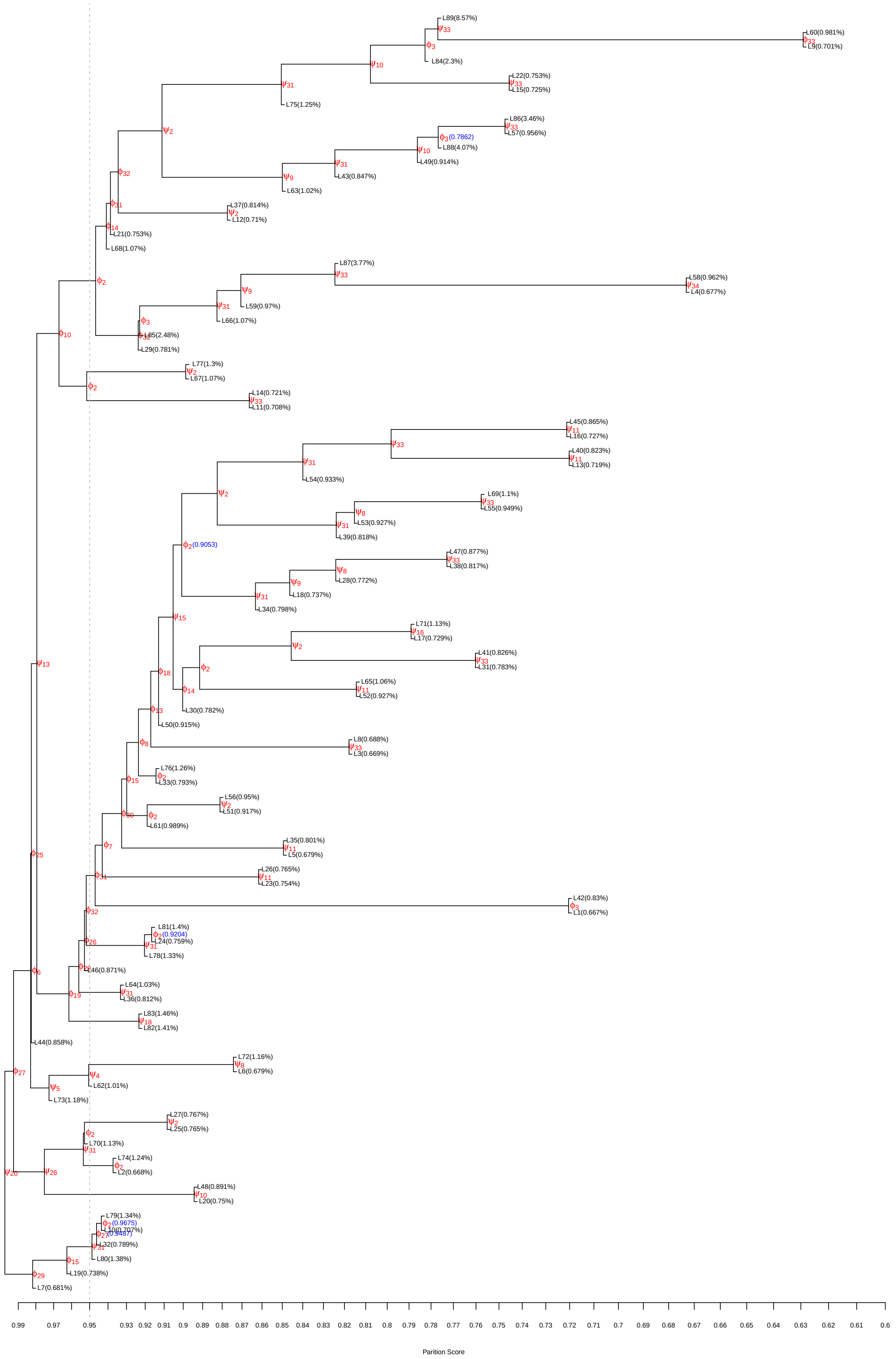

Fig. 14. Partition tree of HP35 Nle/Nle with $S_0 = 10000$, $P_c = P_0 = 0.6$, and the Gaussian kernel.

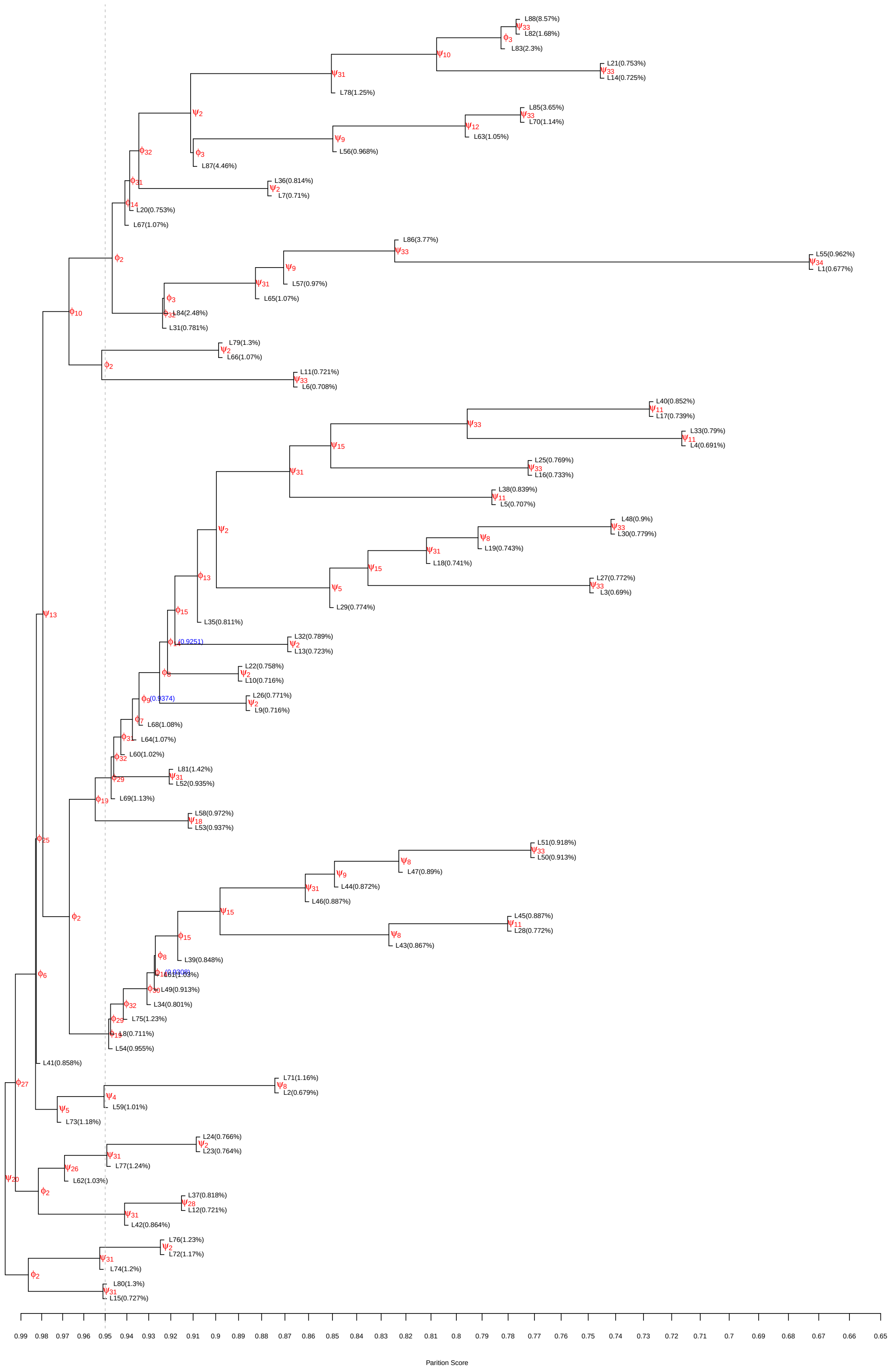

Fig. 15. Partition tree of HP35 Nle/Nle with $S_0 = 10000$, $P_c = P_0 = 0.65$, and the Gaussian kernel.

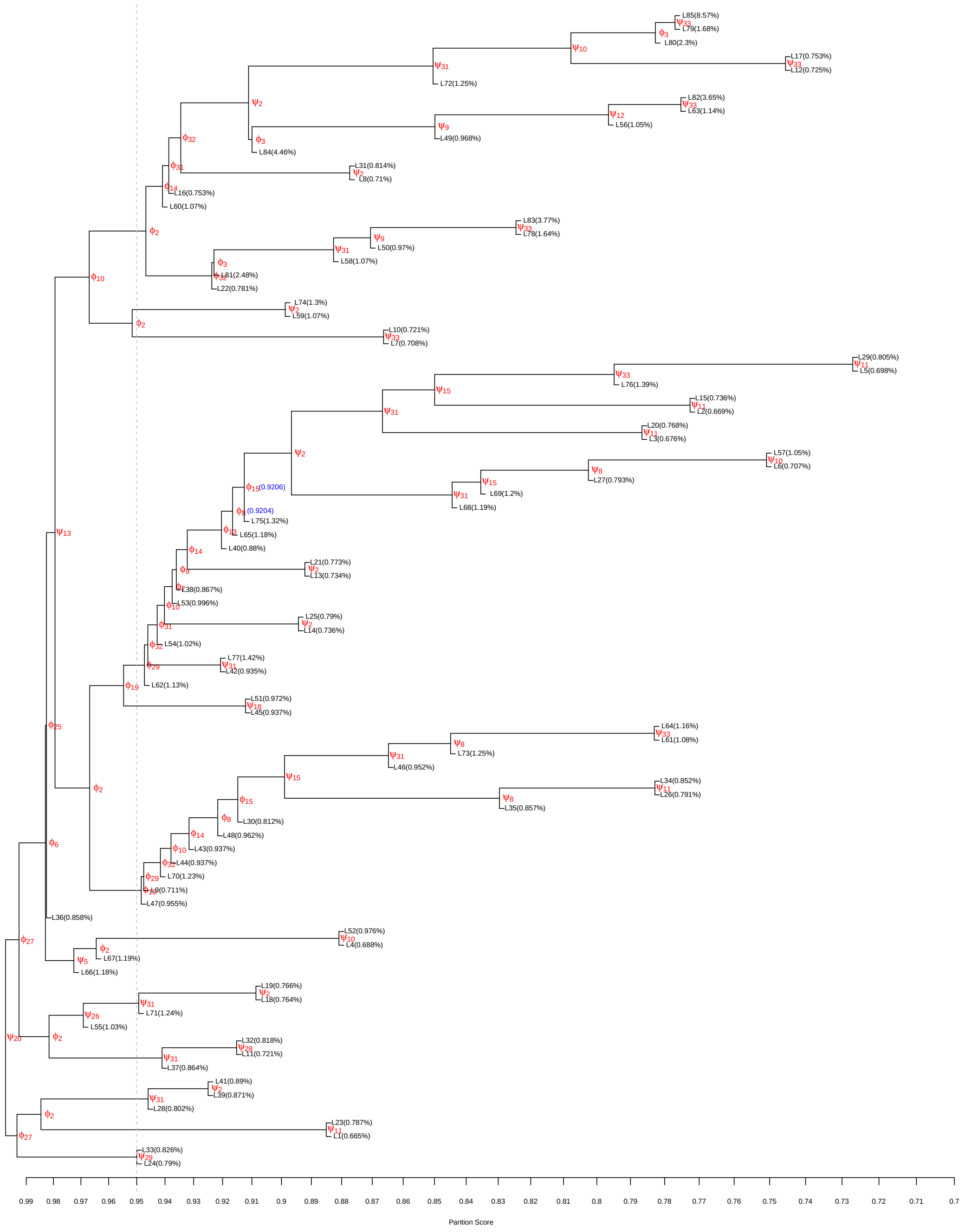

Fig. 16. Partition tree of HP35 Nle/Nle with $S_0 = 10000$, $P_c = P_0 = 0.7$, and the Gaussian kernel.

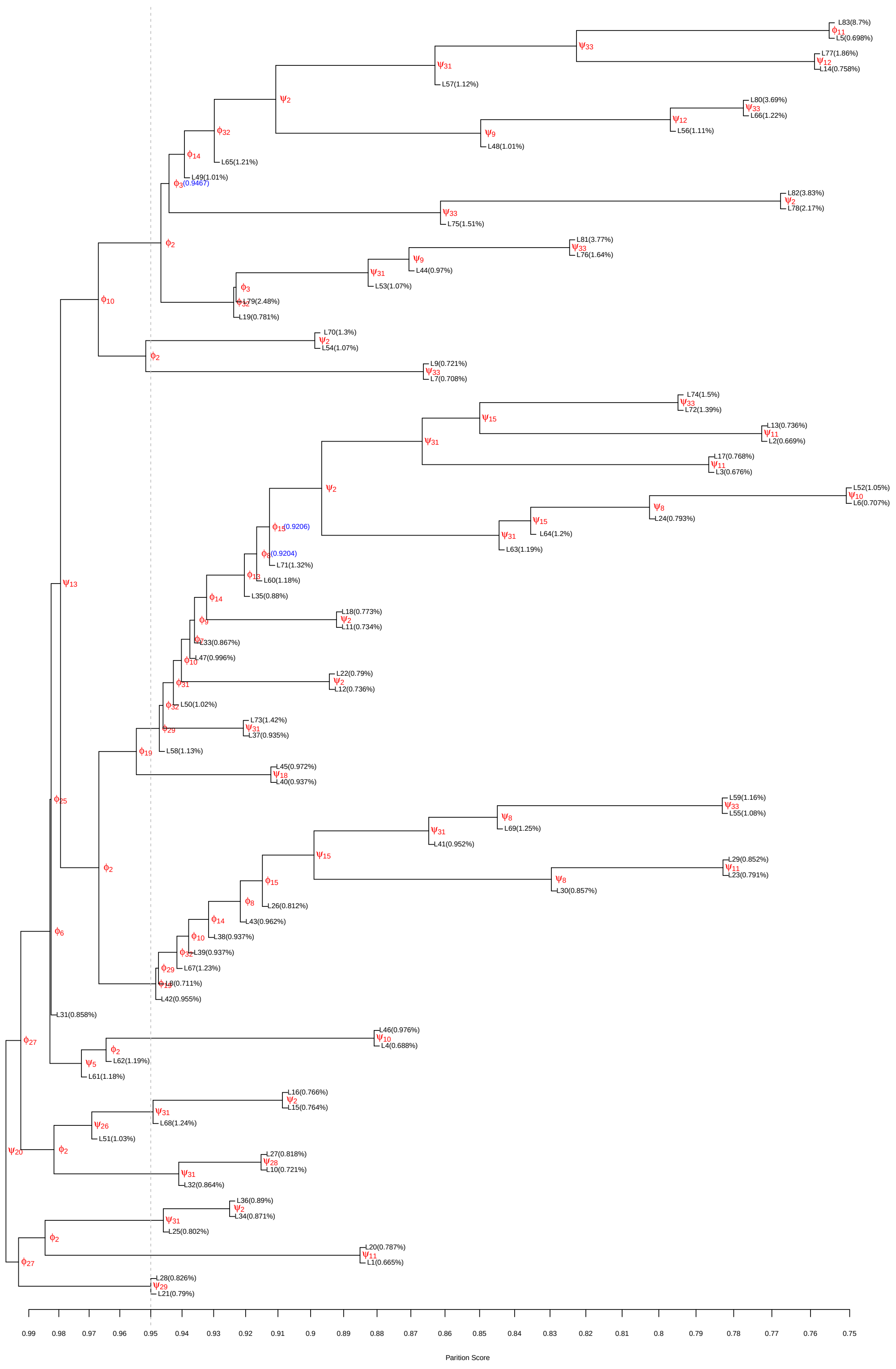


Fig. 17. Partition tree of HP35 Nle/Nle with $S_0 = 10000$, $P_c = P_0 = 0.75$, and the Gaussian kernel.

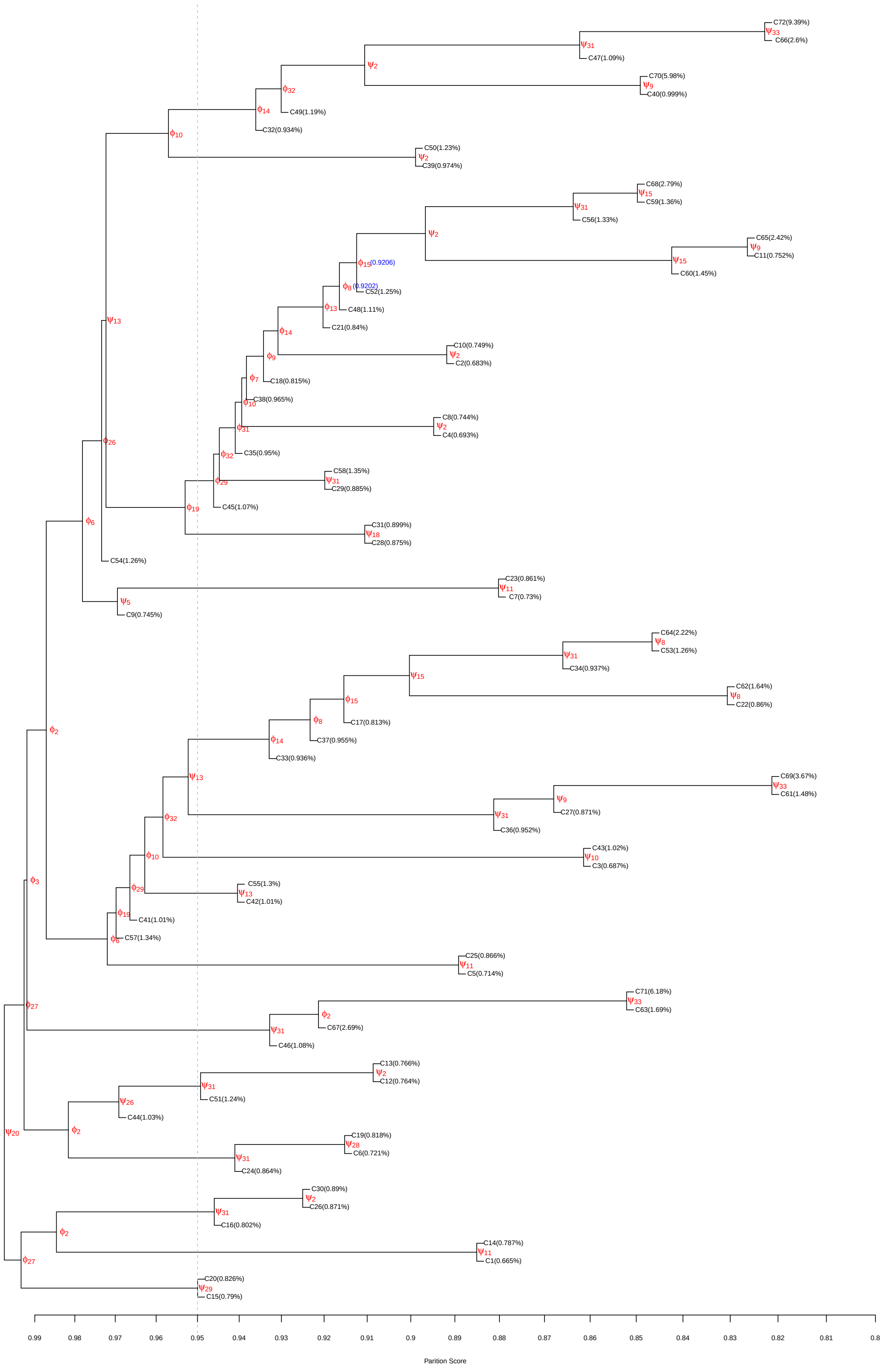


Fig. 18. Partition tree of HP35 Nle/Nle with $S_0 = 10000$, $P_c = P_0 = 0.8$, and the Gaussian kernel.

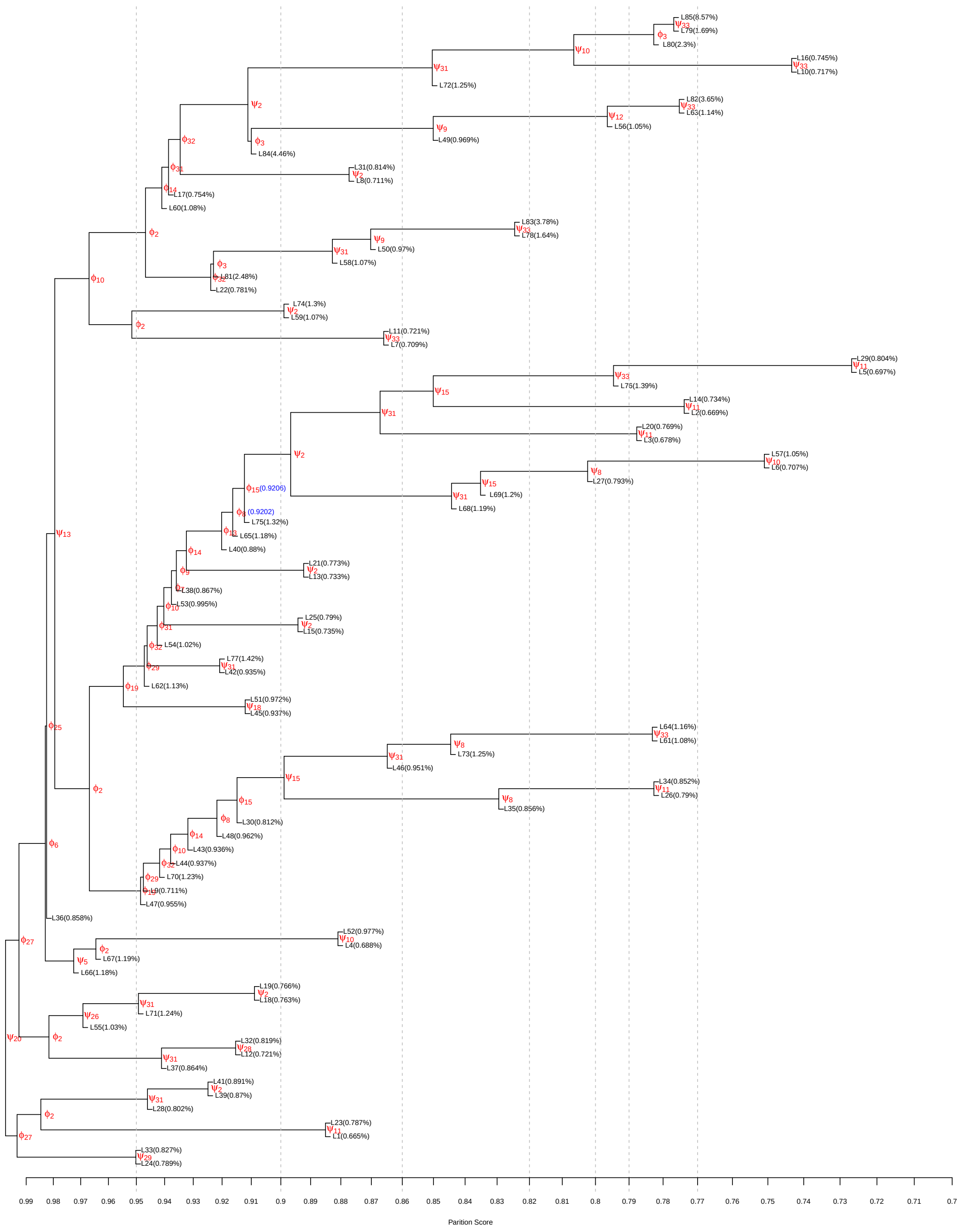

Fig. 19. Partition tree of HP35 Nle/Nle with $S_0 = 10000$, $P_c = P_0 = 0.7$, and the Epanechnikov kernel.

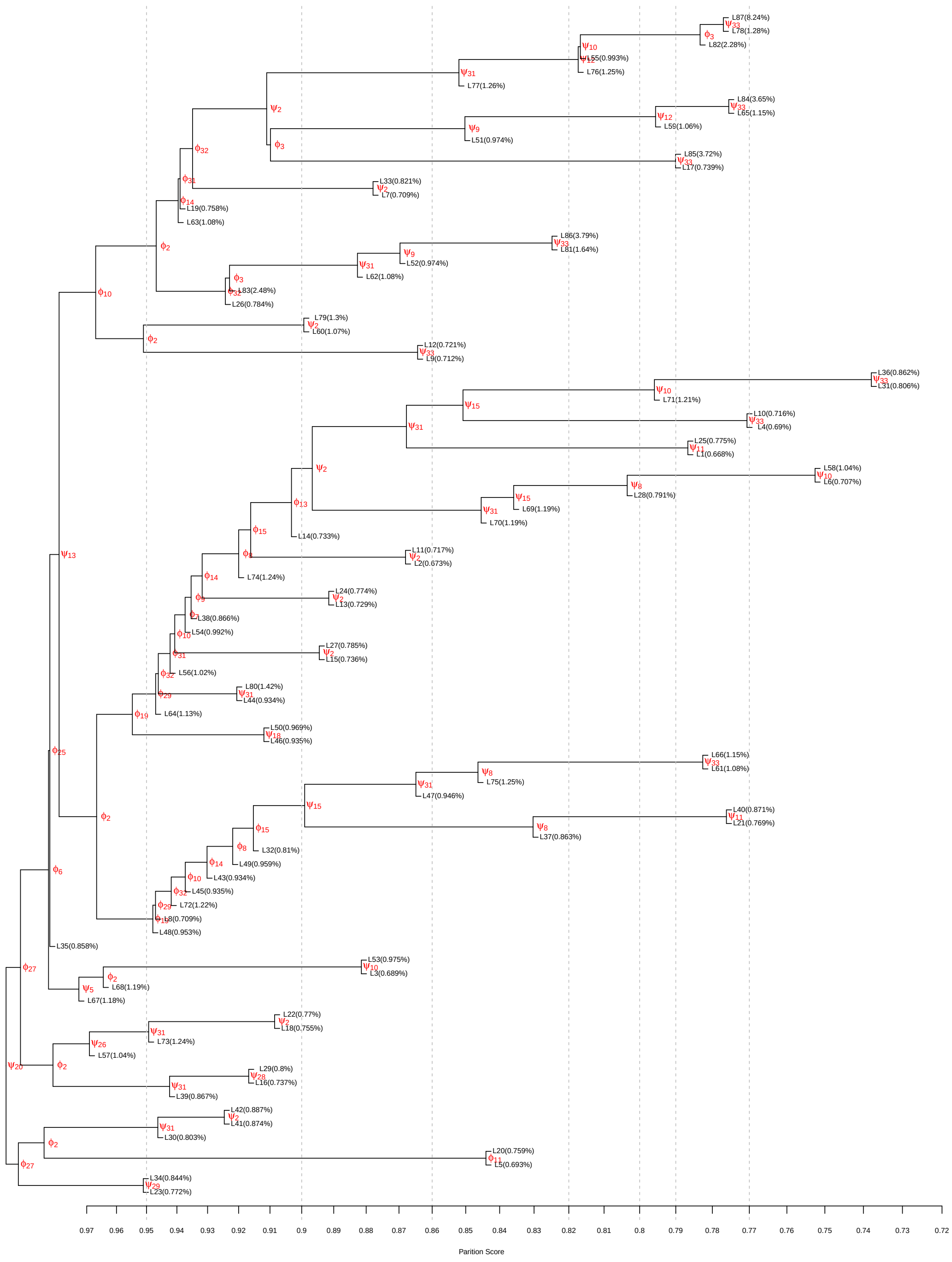

Fig. 20. Partition tree of HP35 Nle/Nle with $S_0 = 10000$, $P_c = P_0 = 0.7$, and the von Mises kernel.
